# Auditory bullae enlarge along an aridity gradient in a diverse subfamily of small carnivorous mammals

**DOI:** 10.64898/2026.11.17.767213

**Authors:** Cameron S. Dodd, Vera Weisbecker, Andrew M. Baker, Meg L. Martin, Kenny J. Travouillon, Linette S. Umbrello, Michael Westerman, Renee A. Catullo

**Affiliations:** School of Biological Sciences, University of Western Australia, 35 Stirling Highway, Crawley, WA 6009 Australia; Collections and Research, Western Australian Museum, 49 Kew Street, Welshpool, WA 6106 Australia; College of Science and Engineering, Flinders University GPO Box 2100, Adelaide, SA 5001 Australia; Australian Research Council Centre of Excellence for Australian Biodiversity and Heritage Wollongong, NSW 2522 Australia; School of Biology and Environmental Science, Queensland University of Technology, 2 George Street, Brisbane, Qld 4001 Australia; Biodiversity and Geosciences Program, Queensland Museum, South Brisbane, Qld 4101 Australia; Department of Environment and Genetics, La Trobe University, Bundoora, VIC 3086 Australia

**Keywords:** Dasyuridae, bullar hypertrophy, sensory ecology, cranial modularity, 3D GMM, generalized dissimilarity modelling

## Abstract

The world’s arid regions are home to diverse assemblages of small mammals, many of which possess a range of physiological, behavioral, and sensory adaptations to the harsh selective pressures imposed by arid environments. The mammalian skull facilitates a range of sensory functions, meaning that arid-dwelling species likely possess distinct skull adaptations which have evolved in response to the unique selective pressures imposed by arid environments. Studies on arid-adapted rodents have highlighted hearing as being important for arid adaptation; however, the adaptive potential of the entire skull is rarely considered. Here, we used 3D geometric morphometrics to test for signatures of arid adaptation using 168 skulls across 30 species of the Sminthopsinae, a subfamily of small insectivorous marsupials, found across Australia. We found that relative bulla size increased by 20% along an aridity gradient from mesic to hyper arid. We also found that species further from the equator tended to have wider posterior nasal cavities, which may be related to selection for thermoregulatory capabilities. Our findings demonstrate the role of the skull in facilitating adaptation to aridity and provide further evidence that selection for improved low-frequency hearing appears to be a key driver of skull shape in arid-zone small mammals.

## Introduction

Arid environments are characterized by low and/or sporadic rainfall, extreme temperatures, and extreme variation in temperature on a daily and seasonal scale (Nash et al., 1999). These conditions generally lead to ecosystems where vegetation is sparse, and organisms must cope with prolonged periods of drought (Dickman et al., 2011; Letnic & Dickman, 2010). Despite these challenges, most arid regions contain a diverse assemblage of small mammals (Kelt, 2011; Kelt et al., 1996), within which a range of physiological, behavioral and sensory adaptations have evolved in response to selection imposed by aridity (Willmer et al., 2000). Many common physiological and behavioral adaptations to aridity, such as improved kidney function (Bankir & de Rouffignac, 1985; Chalopin et al., 2026) and the ability to enter torpor (Levy et al., 2011; Warnecke et al., 2008), are directly driven by abiotic factors such as water conservation (Van Sant et al., 2012) and heat regulation (Bakken, 1976). However, many sensory adaptations, such as sight and hearing, are primarily driven by the biotic sensory environments that arid ecosystems promote i.e. arid ecosystems tend to have reduced vegetation cover and structure (Smith et al., 1990), and low prey/forage availability (Brown et al., 1979; Crawford & Gosz, 1982), which leads to a more physically open environment. As a result, arid environments impose selection for both physiological processes, like efficient heat and water regulation (Willmer et al., 2000), as well as improved long-distance sensory capabilities (Fleischer, 1978; Willmer et al., 2000).

The mammalian skull is important for a range of sensory and physiological functions including sight, smell, feeding and hearing, all of which may face selection to maintain/improve function under challenging arid conditions (Nummela et al., 2013; Smith & Ruf, 2026). Adaptations of the nasal cavity to arid environments may involve increased respiratory turbinal complexity, which may reduce respiratory water loss (Schmidt-Nielsen et al., 1970; Van Sant et al., 2012; Walsberg, 2000). There is some evidence that narrowing nasal cavities may improve the efficiency of respiratory water reabsorption by increasing the respiratory surface area to volume ratio (Inthavong et al., 2009). More complex turbinals (Baldo et al., 2026; Cortés et al., 1990; Nelson et al., 2007; Ruelas et al., 2025) and narrower nasal passages (Alhajeri & Steppan, 2018; Cortés et al., 1990) have been found in arid-zone rodent species, however the turbinals remain highly understudied in general (Martinez et al., 2024), and their dual role in respiration and olfaction (Martinez et al., 2020; Taylor et al., 2024) makes inferring the selective pressures driving changes in nasal structure challenging.

Increased nocturnality in arid mammals, as a mechanism to avoid extreme heat, may lead to changes in the shape and positioning of the orbits (Bennie et al., 2014). On average, nocturnal mammals have larger eyes (Hall et al., 2012) than diurnal mammals, and nocturnal rodents tend to have more forward-facing eyes (improved binocular vision) (Vega-Zuniga et al., 2013). However, eye evolution is complicated by the likely ancestral nocturnality of most mammal groups (Gerkema et al., 2013; Walls, 1942), with many diurnal mammals retaining traits associated with their ancestral nocturnality (Hall et al., 2012). The diets of arid-zone herbivores tend to be tougher and more fibrous, largely due to adaptations for desiccation resistance in many plants (Lee, 2018), and this has been suggested to select for associated changes in jaw shape and chewing musculature (Mitchell et al., 2018; Quibod et al., 2023; Randall et al., 2025; Weisbecker et al., 2019). A similar pattern has also been proposed in insectivores where tougher, desiccation-resistant invertebrate cuticles in arid areas (Fisher & Dickman, 1993b) may lead to changes in jaw shape (Sanson, 1985). However, no clear evidence for this has been found (Fisher & Dickman, 1993b, 1993a). While we may expect mastication apparatus to be more robust in arid species, this region of the skull can be highly plastic during development, and influenced by strong individual-level variation in feeding habits (Cooper et al., 2023; Scicluna et al., 2025; Weisbecker et al., 2019). As a result, it is important to consider the different scales of variation, specifically individual-vs. species-level, as the processes which shape these can be quite distinct (Sherratt & Kraatz, 2023).

The most studied arid skull adaptations in mammals are associated with hearing (Alhajeri & Steppan, 2018; Mason, 2016a, 2016b). Research has largely focused on the prominent hypertrophic bullae (enlarged bony inner ear) present in several arid-zone rodent groups including the Gerbilinae (Alhajeri et al., 2015; Alhajeri & Steppan, 2018; Lay, 1972; Nengovhela et al., 2019; Plassmann & Kadel, 1991; Ravicz & Rosowski, 1997), Murinae (Alhajeri & Steppan, 2018; Edwards & Bessinger, 2024; Nengovhela et al., 2019; Tabatabaei Yazdi et al., 2015), Dipodomyinae (Alhajeri & Steppan, 2018; Lay, 1972; Webster, 1962; Webster & Webster, 1975, 1977; West, 1985), and Dipodidae (Alhajeri & Steppan, 2018; Mason, 2016a). An increase in bulla volume is associated with increased ability to detect low-frequency sound (Ravicz & Rosowski, 1997; Webster, 1962). This is because the bulla acts as a Helmholtz resonator (Plassmann & Kadel, 1991), with a volume inversely proportional to its resonant frequency; thus, a larger bulla allows for greater amplification of lower frequency sounds (Mason, 2016b). Low-frequency hearing is expected to be selected for in arid species as low-frequency sounds travel further, and long-distance hearing is more advantageous in open arid environments than in more densely vegetated areas (Ekdale, 2016; Fleischer, 1978; Mason, 2016a, 2016b).

This evolution of larger bullae in arid-zone species has been identified in several non-rodent groups; for example, in sengis (Macroscelididae) (Benoit et al., 2013; Mason, 2016a), sand cats (*Felis margarita* Loche 1858) (Huang et al., 2002), and in marsupial bandicoots and bilbies (Peramelemorphia) (Taylor et al., 2022). The evolution of larger bullae in arid species has also been suggested in the Sminthopsinae (Archer, 1981), a subfamily of small (mostly < 50g), largely insectivorous marsupials in the family Dasyuridae which consists of four major clades (Krajewski et al., 1997, 2012; Westerman et al., 2016, 2026). The diversification of the sminthopsins has been closely linked with the aridification of Australia through the late Miocene (Black et al., 2012; García-Navas et al., 2018). Multiple lineages within the Sminthopsinae independently colonized the arid zone (García-Navas & Westerman, 2018), suggesting multiple evolutionary transitions between arid and mesic biomes. Archer (1981) speculated that a range of traits may play a role in arid adaptation in the Sminthopsinae, including relative tooth row proportions, bulla size, palatal vacuity length and posterior palate width. Subsequent advances in our understanding of the phylogenetic relationships between sminthopsin taxa (Krajewski et al., 1997, 2012; Westerman et al., 2016, 2026), the description of several new species (AMTC, 2026), and the development of additional statistical tools, means that these hypotheses can now be rigorously tested.

Testing hypotheses regarding the evolution of sensory structures in the mammalian skull across taxa can be challenging, as the bones of the skull do not vary independently and may be bound by a complex network of covariance relationships (Goswami, 2006; Novacek, 1993). This can be due to underlying developmental covariance between structures (Lande, 1986), functional interactions across the skull (Felice et al., 2018), complex selective pressures acting across multiple structures at once (Marcy et al., 2024; Morrissey, 2014), and the potential for substantial developmental plasticity (Weisbecker et al., 2019). Each of these factors can be species specific, and skull evolution can be further constrained by allometry (size-associated effects) (Marcy et al., 2020, 2024; Mitchell et al., 2024) and phylogenetic signal (Mitchell et al., 2026). Understanding how each of these factors can constrain and facilitate changes in skull shape is crucial for understanding adaptation across different skull structures, and between taxa.

Here, we used a 3D geometric morphometric approach to investigate how patterns of cranial variation across the Sminthopsinae may have facilitated arid adaptation. We assessed patterns of covariance across the skull using the modularity framework (Goswami, 2006; Zelditch & Goswami, 2021), which defines discrete ‘modules’ across the skull that capture the strongest patterns of covariance between landmarks. Using this framework, we also investigated whether patterns of module disparity (breadth of shapes observed) and integration (covariance of landmarks) correlated with patterns of arid adaptation. Additionally, we investigated the role of allometry (shape differences related to size) and phylogenetic signal in skull shape evolution. We expected that variation in the bulla shape would be highly correlated with aridity, as has been found in other small mammals (Alhajeri & Steppan, 2018; Mason, 2016a; Taylor et al., 2022), but that correlation with environment may be detectable across other functional skull regions.

## Materials and Methods

### Sampling and 3D scanning

We sampled a total of 168 crania from all 25 currently recognized species within the Sminthopsinae, as well as five lineages not formally recognized, but proposed to represent distinct species (Blacket et al., 2001; Cooper et al., 2000; Dodd et al., 2025). Skulls were obtained from the Australian Museum Sydney (AMS), Museum and Art Gallery of the Northern Territory (MAGNT), Museums Victoria (MV), Queensland Museum (QM), South Australian Museum and Art Gallery (SAMA), and Western Australian Museum (WAM) (Supplementary S1). We digitized all skulls using X-ray microcomputed tomography (*μ*CT) scan data with a Nikon XT H225 ST CT-scanner at the Australian Museum Research Institute (AMRI) (all skulls from the AMS) or the same model of scanner at the Western Australia National Imagine Facility (NIF) node at the Centre for Microscopy, Characterisation and Analysis (CMCA) at the University of Western Australia (UWA) (all other skulls). Scanning parameters were as follows: AMS, scan=circular, voltage=180kV, current=61uA, gain=24dB, exposure=250ms; CMCA, scan=helical, voltage=75kV, current=75uA, gain=24dB, exposure=2000ms. We reconstructed X-ray images into cross-sectional image stacks using CT Pro 3D (Nikon Metrology, Tring, UK), segmented out artefacts and built 3D meshes using CTAnalyser v1.20.8.0 (Bruker, 2020), and finally decimated meshes to ∼500,000 faces using MeshLab v2025.07 (Callieri et al., 2012) to aid in file size management.

### Phylogeny

To test the effect of phylogenetic relationships on skull shape, we used the phylogeny from Kealy and Beck (2017). This phylogeny is consistent with other studies (Krajewski et al., 1997, 2012; Westerman et al., 2016, 2026), which show that the Sminthopsinae consists of four monophyletic clades: *Sminthopsis macroura* clade, *Sminthopsis murina* clade*, Ningaui* and *Antechinomys*. We trimmed the phylogeny to include only the Sminthopsinae using *phytools::keep.tip()* (Revell, 2024) and grafted species not represented onto the phylogeny using *phytools::bind.tip()* at a position of half the terminal branch length. The phylogenetic placement of *Antechinomys spenceri* (Thomas, 1906), *Antechinomys auritus* Dodd, Catullo, Travouillon, Baker, Westerman and Umbrello, 2025, and *Antechinomys* sp. nov.‘south-west Western Australia’ followed Dodd et al. (2025). The position of *Sminthopsis macroura* (Gould, 1845) ‘South clade’ and *S. macroura* ‘West clade’ followed Blacket et al. (2001). The position of *Sminthopsis nitela* (Collet, 1897) followed Umbrello et al. (2024).

### Landmarking

The lead author landmarked all skulls using Checkpoint (Stratovan, 2025) to avoid operator bias (Fruciano et al., 2017).We anonymized the meshed by assigning a random ID number to each and landmarked them in that random order. For each mesh, we initially placed 40 anchor landmarks and then projected the others using the template function in Checkpoint. We then manually adjusted each landmark in a non-standardized order. We discarded and reprocessed the first 20 sets of landmark data to reduce error during template familiarization, and we landmarked a further 26 crania twice to quantify landmarking repeatability (Marcy et al., 2020), giving a total of 194 datasets from 168 samples. In total, we placed 98 point-landmarks and 37 curves on each mesh, totaling 274 landmarks. We adapted the landmarking protocol from Randall et al. (2025), and for full landmark descriptions see Supplementary S2. We did not landmark the zygomatic arches given the potential for them to warp or break during specimen preparation (Schmidt et al., 2010; Yezerinac et al., 1992); however, we did landmark the points where the arches meet the skull. We imported the landmarks into R (R Core Team, 2025) where we carried out all further data manipulation and analyses were performed, largely using *geomorph.v4.1.10* (Adams et al., 2025). We will use the notation *package::function()* to detail the key R packages and functions used following the package’s first mention. All code required to replicate landmark data manipulation, analyses and plots are available on GitHub.

### Landmark processing

We imputed missing landmark data within species using *geomorph::estimate.missing()*, with the maximum missing data being 12%. For species with less than five skulls, we imputed missing data for each skull individually, using the ten most similar skulls by Procrustes distance. We aligned landmark coordinates using Generalized Procrustes Analysis (GPA) (Rohlf & Slice, 1990) using *geomorph::bilat.symmetry* (Baken et al., 2021), taking cranial symmetry into account (Klingenberg et al., 2002) and using replicate samples to quantify landmarking error. We optimized semilandmark positions by sliding during landmarking, so semilandmarks were not allowed to slide during GPA analyses. There is debate as to the impact of using a single whole-skull GPA and then subsetting landmarks into modules, or performing separate, module-specific GPAs (Baab, 2013; Cardini, 2019; Klingenberg, 2009). We primarily used module-specific GPAs so that variance is scaled relative to module size. This meant that our modules no longer existed in the same Procrustes space. However, we replicated all analyses using a single whole-skull GPA (see Supplementary Material) to assess whether this has a meaningful effect on the interpretation of our results.

### Modularity hypothesis testing

To assess variation in shape across different skull regions, we first had to split the skull into separate regions (modules) on which we could model evolution. We used the modularity framework (Cheverud, 1982; Goswami, 2006; Zelditch & Goswami, 2021) to define discrete modules that best described patterns of landmark covariance across our dataset. We generated several candidate modularity hypotheses based on existing literature (Cheverud, 1982; Goswami, 2006; Willmore et al., 2006; Zelditch, 1988) and tested the best fit to our data using *EMMLiv2::subsampleEMMLi()* (Goswami & Finarelli, 2016). This method uses a maximum likelihood approach to test which hypothesis best fits the covariance patterns of a set of landmarks. Full methods are detailed in the Supplementary Methods, all candidate hypotheses are shown in Supplementary S3 and a summary of the comparisons between hypotheses is shown in Supplementary S4. In short, our dataset was best described using a six-module system consisting of basicranium, bulla, molar, orbital, rostrum, and vault (Figure 1). This is largely consistent with other studies on mammal skulls (Cheverud, 1982; Goswami, 2006; Heck et al., 2018; Randau et al., 2019; Sherratt & Kraatz, 2023), however unlike these previous studies, we found that the basicranium was best described as two modules: the bulla and the rest of the basicranium (Figure 1).

**Figure 1.**
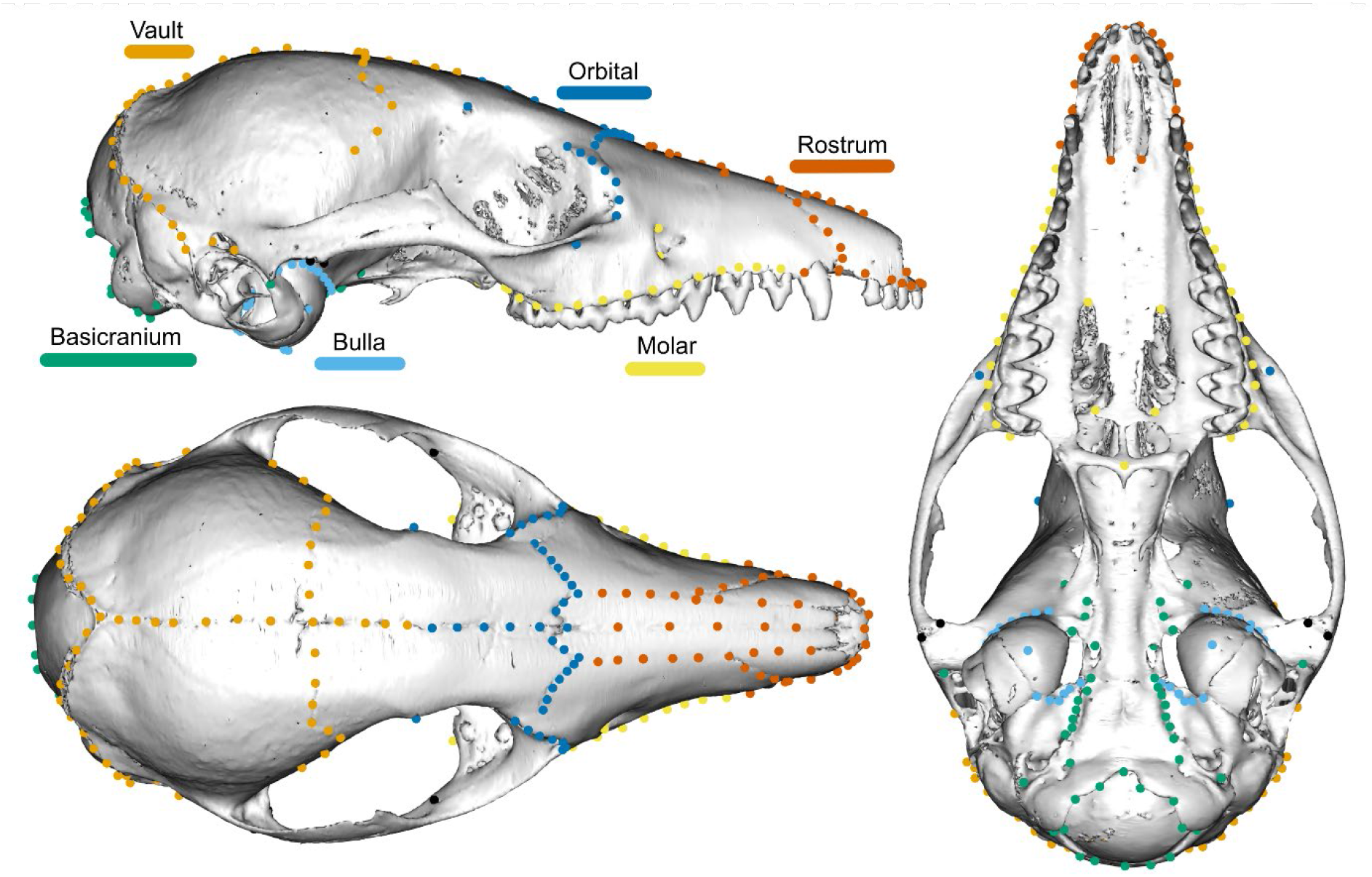
Cranial landmarks used in this study. Landmarks are colored according to modularity hypothesis adopted by this study (see Supplementary Methods), with black landmarks not belonging to any module. Full landmark descriptions can be found in Supplementary S2. The mesh is the skull of a *Sminthopsis granulipes* (Troughton, 1932) (WAM M17501).

### Allometry and phylogenetic effects

We tested the amount of shape variation attributable to cranial allometry using *geomorph::procD.lm()* following a single whole-skull GPA and using skull centroid size as our size metric. We tested the effect of allometry with and without an interaction between clade and size to examine the potential for different allometric slopes between genera. We examined shape variation independent of phylogeny using a phylogenetic Principal Component Analyses (phyloPCA) (Revell, 2009), implemented in *geomorph::gm.prcomp()* for the whole skull and each module individually. We used the Köppen-Geiger climate classification region to define three biomes across Australia: arid, climate types *BWh (Hot desert)* and *BWk (Cold desert)*; semi-arid, climate types *BSh (Hot semi-arid)* and *BSk (Cold semi-arid)*; and mesic, all other climate types in Australia (Figure 2). Species were assigned biomes based on expert distribution maps from Baker and Gynther (2023) and occurrence data downloaded from the Atlas of Living Australia (ALA), with primary literature being used for species whose taxonomy has not yet been updated in these sources (Blacket et al., 2001; S. J. B. Cooper et al., 2000; Dodd et al., 2025; Umbrello et al., 2024). We then investigated the relative disparity of each module using *geomorph::morphol.disparity()*, the between-module integration using *geomorph::two.b.pls()* and the within-module integration using *geomorph::integration.Vrel()*. We also calculated the evolution rate of the shape of each module, assuming Brownian Motion, using *geomorph::compare.multi.evol.rates()* (Denton & Adams, 2015).

**Figure 2.**
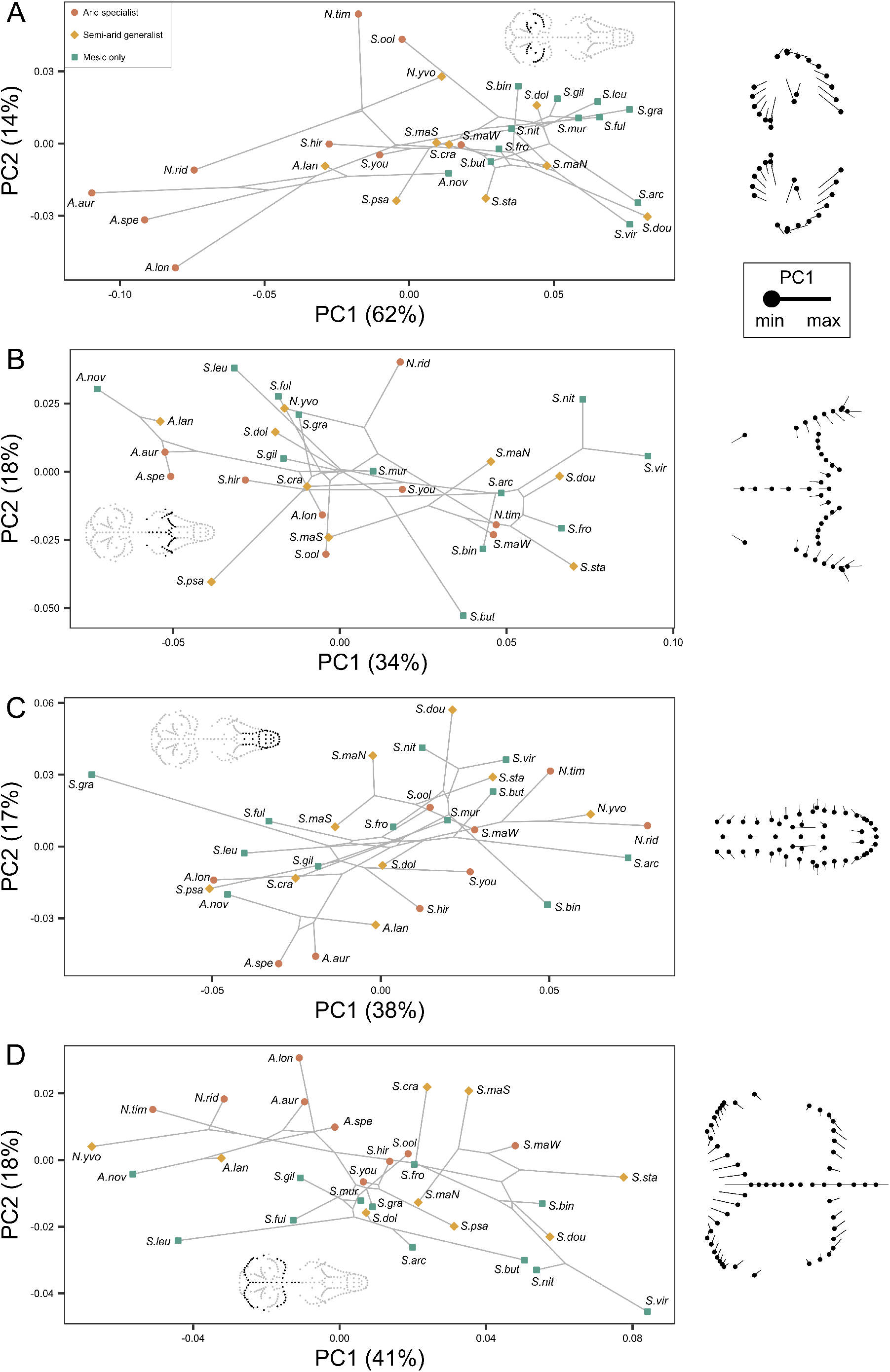
Phylogenetic PCA (Revell, 2009) of skull shape within the Sminthopsinae by module: A) bulla, B), orbital, C) rostrum, D) vault. Species are colored by environment. Analyses were carried out using the mean shape for each species following separate module-specific GPAs. Displacement vector plots show the changes in shape from the minimum observed PC1 value (solid circle) to the maximum observed PC1 value (end of vector) for each module. Plots of whole-skull shape, basicranium and molar modules can be found in Supplementary S6. Similar plots for all modules generated using a single whole-skull GPA and allometry corrected data can be found in Supplementary S7-8 respectively.

### Arid adaptation

To test how skull shape changes with increasing aridity, we tested associations between module shape and environment using generalized dissimilarity modelling (GDM) (Ferrier et al., 2007). GDM predicts trait (module shape) dissimilarity from environmental differences between sites (samples), and has the advantage of using I-spline functions to detect non-linear relationships and variation in the rate of phenotypic change along environmental gradients (Mokany et al., 2022). By comparing the associations between environmental predictors and shape dissimilarity across modules, we can identify environmental variables differentially associated with module shape, providing evidence consistent with module-specific responses to environmental selection pressures. We calculated shape dissimilarity among samples separately for each module using Procrustes distance (Baken et al., 2021; Rohlf & Slice, 1990). We obtained environmental predictors from the Worldclim bioclimatic variables (Fick & Hijmans, 2017), with two additional layers representing an aridity index and potential evapotranspiration (Zomer et al., 2022), totaling 21 layers. To reduce collinearity among predictors, we cropped environmental raster layers to the geographic extent of the sample coordinates, and calculated Pearson’s correlation coefficients between variables (Pearson & Lee, 1903). We then removed highly correlated layers until no layer-layer pair had an |*r|* > 0.7, leaving a final predictor dataset of nine layers (Supplementary S5). We included geographic distance (in meters), skull centroid size, and genetic distance as covariates in the GDM. We calculated genetic distance as the cophenetic distance between tips in the species-level phylogeny described above, with within-species samples added as a polytomy to each species tip using *ape::cophenetic.phylo()* (Paradis et al., 2004). We treated within-species genetic distances as negligible and set them to zero branch lengths, using a similar approach to Felsenstein (2008). We chose significant predictors of module dissimilarity by stepwise parameter removal comparing model fit to 1000 random permutations of the environmental variables using *gdm.varImp()* in *gdm* (Fitzpatrick et al., 2025) with a significance threshold of p < 0.01. Generalized dissimilarity modelling only detects associations between predictor dissimilarity and morphological dissimilarity, not the direction of that relationship (positive or negative) (Ferrier et al., 2007; Mokany et al., 2022). To resolve this, following the identification of an association, we carried out a post-hoc linear regression of environment against module shape to determine the direction of the relationship.

## Results

The mean missingness prior to imputation was 0.96%, and only 8/168 samples had > 5% missing data (maximum of 12%). In the whole-skull GPA dataset, 8.0% of the variance in shape was attributable to cranial asymmetry, and landmarking errors across duplicated individuals accounted for 1.7% of the variance, showing very high repeatability in the landmarking protocol. Skull size accounted for 14% (R^2^ = 0.14, p < 0.001) of the shape variance, and 26% of shape variation was attributable to differences between clades (R^2^ = 0.26, p < 0.001).

Given the shape differences between clades, we performed all further analyses either using clade as a covariate or using a phylogenetic correction. We detected a small but significant interaction effect between size and clade (R^2^ = 0.05, p < 0.001), suggesting that the four clades may not share a single common allometric slope. However, given the effect size of this interaction was very small compared to other predictors, as well as differences in sample size and skull size distributions across clades, we were not confident in defining clade-specific allometric slopes from this dataset. Thus, we retained a single allometric slope in our analyses.

For the whole skull, the first two PCs represented 45% of the shape variation (Supplementary S6A), less than each module individually (all > 50%) (Figure 2). The PC1 loading on the bulla module was particularly high (62%), suggesting this axis of variation (which largely corresponded to bulla size) was one of the most integrated (strong internal covariance) patterns of skull shape across the Sminthopsinae. There was also no overlap in PC1:PC2 bulla space between the hyper-arid and mesic species (Figure 2A), consistent with our expectations of the bulla showing evidence of arid adaptation. The orbital module also showed minimal overlap between hyper-arid and mesic species, with only the hyper-arid *Ningaui ridei* Archer 1975 clustering with mesic species (Figure 2B).

Correction for allometry had minimal effects on the relative placements of species across PC1:PC2 space for the whole skull and each module (Supplementary S7), although *Ningaui* (containing the smallest skulls) tended to be less distinct following an allometric correction. Performing a single skull-wide GPA before separating the data into modules only had a major effect on relative placements of species in PC1:PC2 space in the basicranium and vault modules (Supplementary S8). For the basicranium, the relative placements of species in PC1:PC2 space became less similar to the bulla when using a single GPA, and for the vault *Ningaui* became far more distinct along PC2.

The strongest between-module integration was found between adjacent modules, as would be expected due to developmental covariance: basicranium:bulla (r = 0.70), rostrum:orbital (r = 0.68), and molar:rostrum (r = 0.62) (Figure 3A). Correlations were notably weaker between the orbital:molar (r = 0.44) and vault:basicranium (r = 0.52) despite being adjacent (Figure 3A). The orbital module displayed the highest disparity, followed by the rostrum and then all other modules (Figure 3B). The basicranium module was the least integrated, followed by the bulla, and the other modules, which were equally integrated (Figure 3C). The rates of evolution largely reflected the disparity of each module; however, the bulla and basicranium evolved faster than expected under Brownian Motion given their observed disparity (Supplementary S10), potentially indicative of a role in arid adaptation.

**Figure 3.**
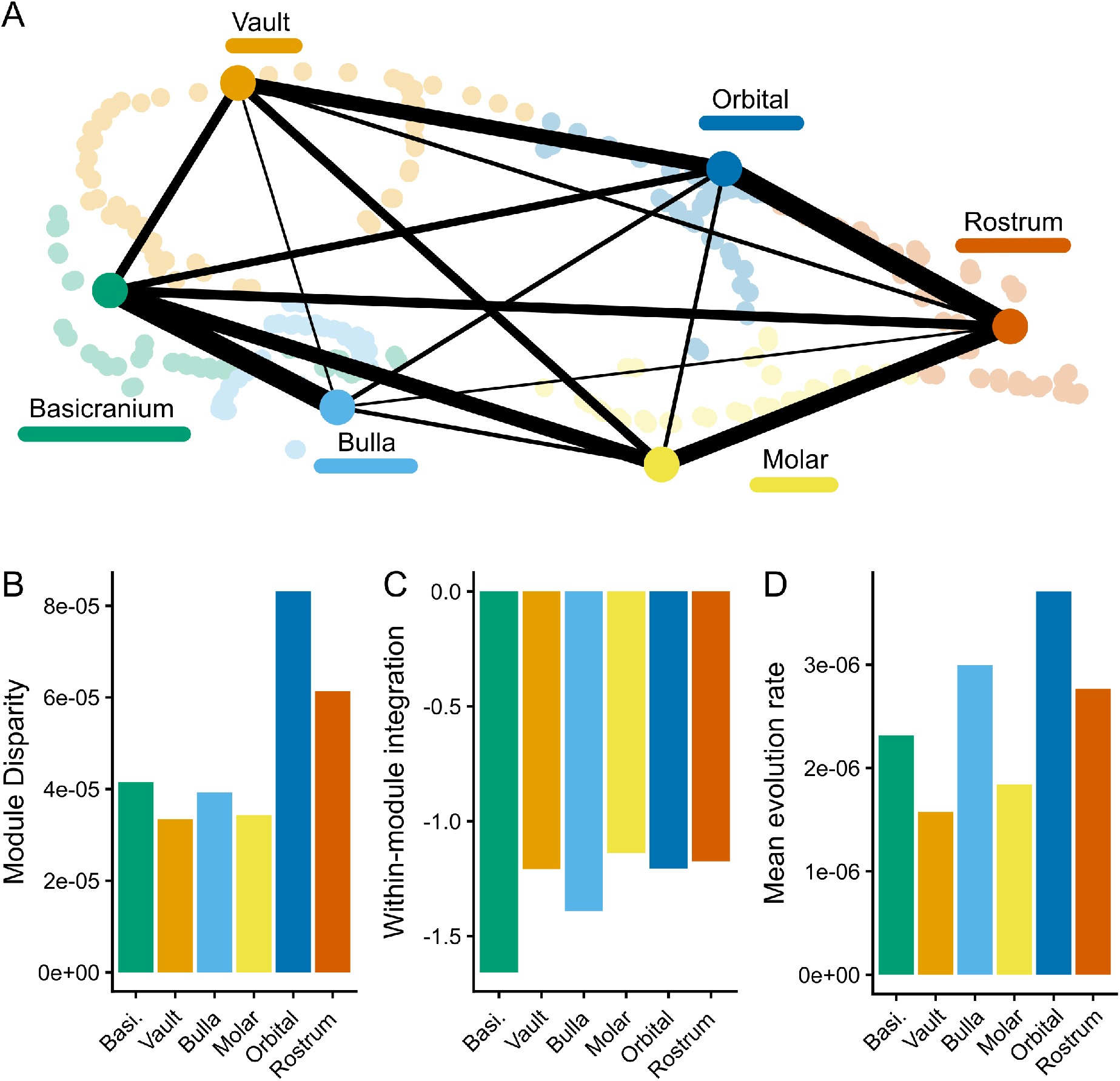
Patterns of disparity and integration between modules across the sminthopsin skull. (A) Relative between-module integration, width of edges represents relative covariance between landmarks of each module pair, r values range from 0.41 (bulla:vault) to 0.70 (bulla:basicranium). (B) Total disparity (Procrustes variance) of each module. (C) Within-module integration, lower (more negative) values indicate weaker integration. Lines joined to bars indicate significant differences between integration values (D) Evolution rate of each module under a Brownian Motion model of evolution. All metrics were calculated following separate generalized Procrustes analyses (GPAs) for each module, meaning they are relative to module size, not absolute size. Similar plots generated using a single whole-skull GPA can be found in Supplementary S9.

For all modules, genetic distance and skull size were significant predictors of morphological dissimilarity, with genetic distance being the only predictor demonstrating a strong association for all modules (Figure 4). Only temperature seasonality and latitude passed an importance of 0.2 for any module; we investigated these with post-hoc analyses of module shape against predictor (Figure 5). As genetic distance and skull size were significant predictors of shape for both modules, we removed the allometric effect of shape and also tested each clade separately.

**Figure 4.**
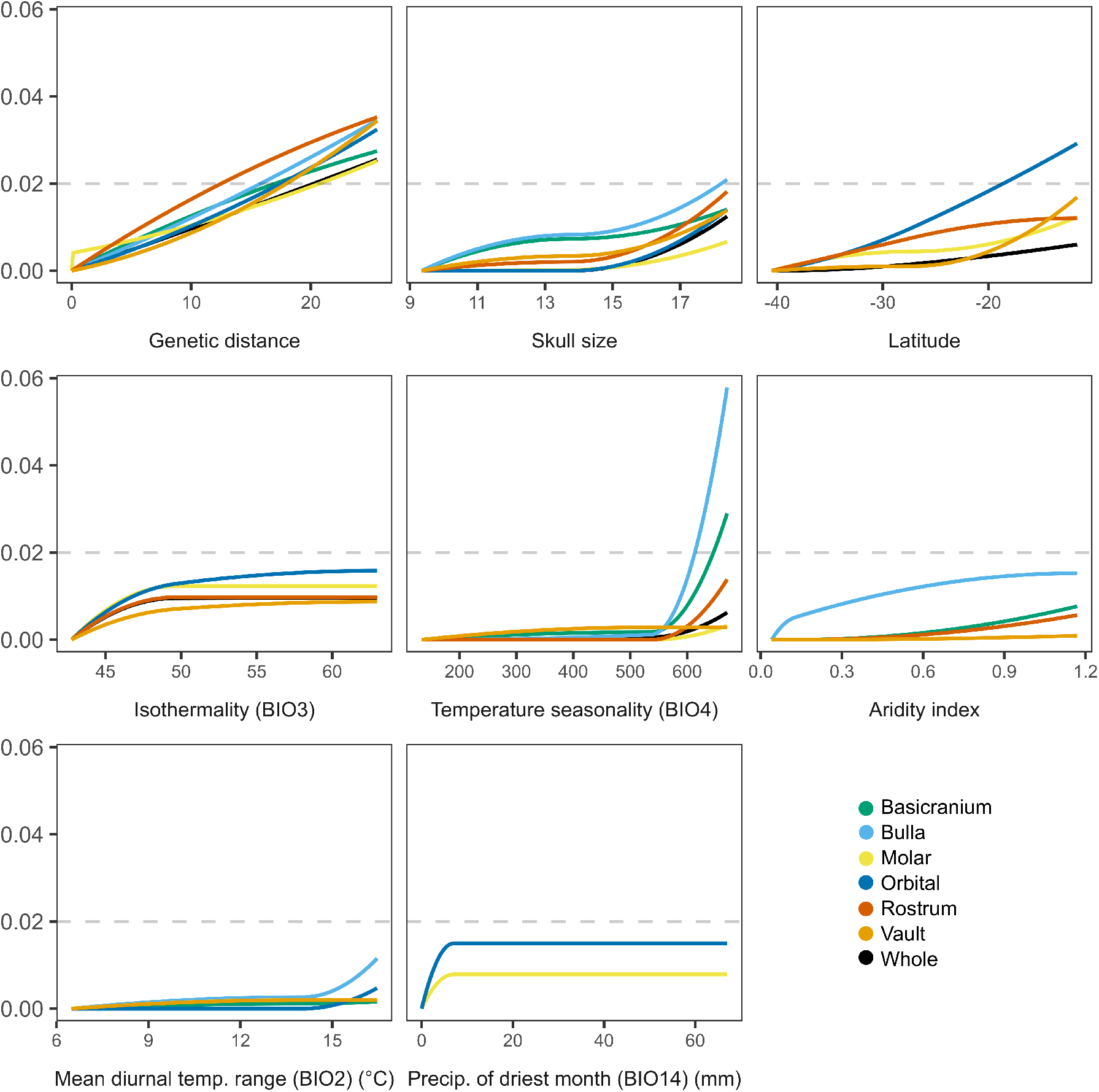
Significant predictors of morphological dissimilarity across the sminthopsin skull following a generalized dissimilarity analysis (GDM). Each module is represented by a different color, the black line corresponds to the pooled whole-skull dataset. The y-axis shows the relative explanatory power of each predictor upon module shape, higher values suggest a stronger association. Predictors which were significant but did not reach an explanatory power of 0.01 for a given module are not shown. Full I-spline parameters can be found in Supplementary S11.

**Figure 5.**
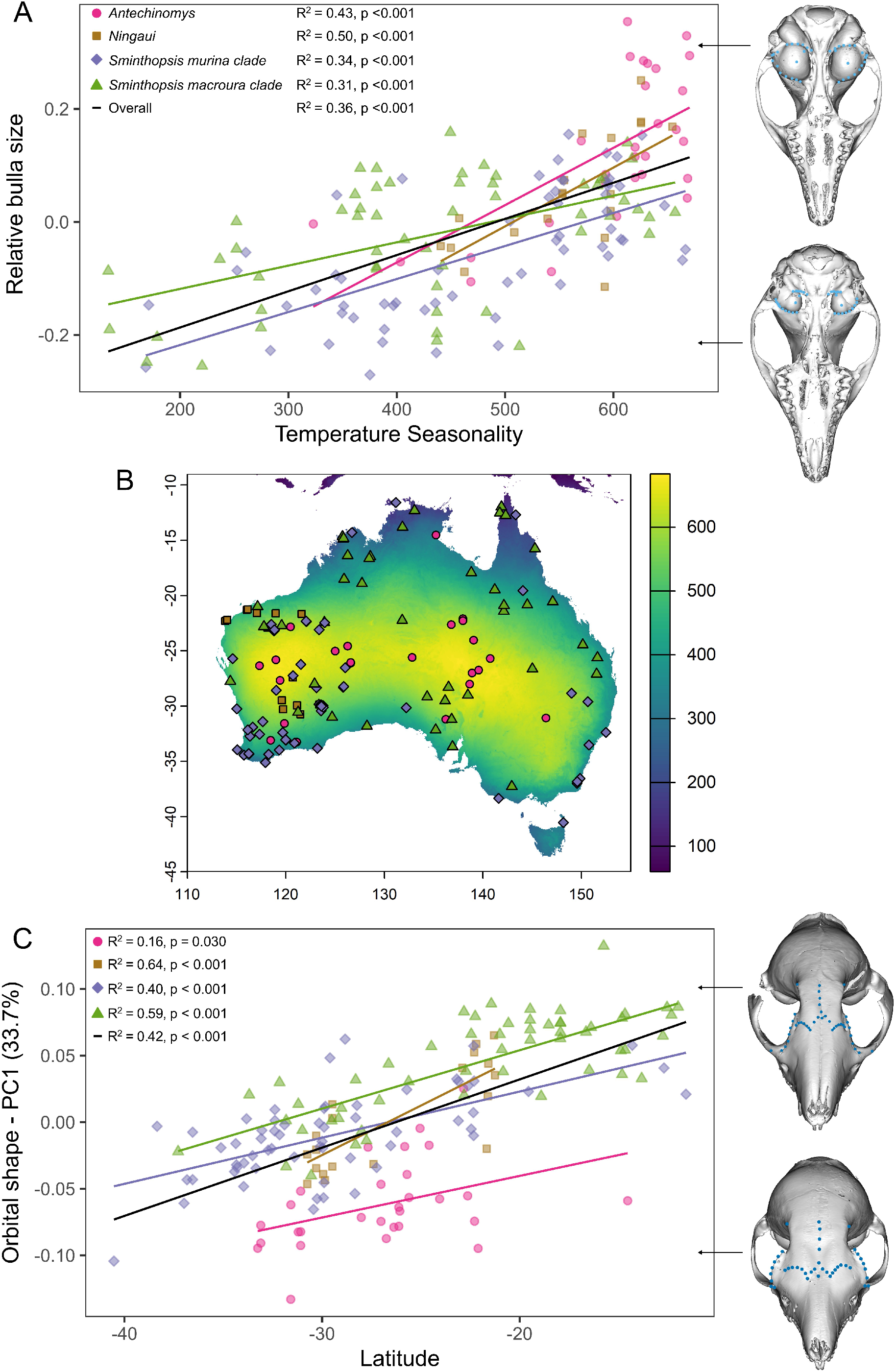
Post-hoc comparisons of the two strongest shape-environment relationships identified following GDM modelling. A) Linear regressions of relative bulla size and temperature seasonality for each clade. Relative bulla size was calculated as the residuals of a regression between bulla centroid size and ‘rest of the skull’ centroid size. Meshes show extreme examples of relative bulla size: large (above) in WAM M56543 (*Antechinomys longicaudatus* (Spencer, 1909)), and small in AM M17302 (*Sminthopsis leucopus* (Gray, 1842)). B) Map of Australia showing the distribution of temperature seasonality and sample localities colored by clade. C) Linear regressions of PC1 of orbital shape against latitude for each clade. Meshes show the shape variation at either extreme of the PC values: WAM M20649 (*Antechinomys* sp. nov.) at the negative extreme and QM JM869 (*Sminthopsis virginiae*) at the positive extreme.

As variation in bulla shape was almost entirely explained by bulla centroid size (ProcD.lm: R^2^ = 0.92, p < 0001), we used relative bulla size to investigate the association with temperature seasonality. We calculated relative bulla size by performing a regression of bulla centroid size against ‘rest-of-skull’ centroid size and extracting the residuals (Figure 5A). For the whole dataset, and each clade individually, we found that increases in temperature seasonality were associated with increases in relative bulla size (Figure 5A). This pattern was also found when samples from the most arid areas (temperature seasonality > 600) were excluded (R^2^ = 0.26, p < 0.001) (Supplementary S12). This suggests that there is a gradient in selection for bulla size along the aridity gradient, and that the relationship identified is not solely driven by samples from the most extreme arid environments.

To test associations between orbital shape and latitude, we used two approaches. First, we performed a PCA of orbital shape on all skulls, and then a linear regression of PC1 with latitude (Figure 5C). We tested all samples as well as each clade individually. Second, we performed a Procrustes ANOVA of latitude against orbital shape (Supplementary S13) on all samples, and separately for each clade. In both approaches, we first used a Procrustes ANOVA to remove shape explained by allometry before testing against latitude. For both approaches, across all genera, we found that as latitude increased (closer to equator), the orbital region became narrower (Figure 5C). This pattern was clearest in *Sminthopsis macroura* clade and *Sminthopsis murina* clade, both due to larger sample sizes and a more even distribution of samples across latitudes.

## Discussion

In this study, we showed strong associations between skull morphology and aridity in sminthopsins, most notably a relative enlargement of the auditory bullae in species from more arid environments. Variation in sminthopsin skull shape was largely structured into six modules corresponding to distinct functional regions. Allometric effects on skull shape were relatively weak, and several previously unrecognized clade-specific patterns of morphological variation were identified. These findings are largely consistent with other studies on arid-adaptation in small mammal skulls (Alhajeri et al., 2015; Alhajeri & Steppan, 2018; Benoit et al., 2013; Edwards & Bessinger, 2024; Huang et al., 2002; Lay, 1972; Mason, 2016a, 2016b; Nengovhela et al., 2019; Plassmann & Kadel, 1991; Ravicz & Rosowski, 1997; Tabatabaei Yazdi et al., 2015; Taylor et al., 2022; Taylor et al., 2024; Webster, 1962; Webster & Webster, 1975, 1977; West, 1985), suggesting that selection for low-frequency hearing, mediated by degree of habitat openness, may be a near universal selective pressure faced by small mammals of all biomes.

We found strong evidence for repeated, convergent evolution of enlarged auditory bullae in response to aridity across the Sminthopsinae, with arid-zone species exhibiting significantly larger bullae than mesic species in each of the four clades. This pattern was retained even when the most arid species were excluded, suggesting that selection for bulla size operates across the aridity gradient, rather than emerging only under extreme conditions. Although hearing studies in the Sminthopsinae are insufficient to determine whether large bullae improve low frequency hearing in this group (Aitkin, 1998; Aitkin et al., 1994; Garrett et al., 2019; Old et al., 2020), the morphological patterns identified here are consistent with findings in rodents, suggesting a similar adaptive mechanism. Experimental reduction in bullar volume has been shown to make rodents *(Dipodomys* and *Meriones*) incapable of avoiding owl (Lay, 1974; Webster, 1962; Webster & Webster, 1971) and rattlesnake strikes (Webster, 1962; Webster & Webster, 1971), both of which are characterized by a burst of low frequency sounds upon strike (Webster, 1962). As a result, bulla size may be driven by a general selection for improved long-distance hearing in open arid environments, including selection for improved prey detection and intraspecific communication; as well as selection specifically for detecting strikes by owls and snakes as key predators in arid ecosystems (McDowell & Medlin, 2009; Morton & Martin, 1979).

While bulla size and aridity were correlated in all sminthopsin clades, only the *Antechinomys* exhibit levels of bullar hypertrophy comparable with that seen in rodents (Alhajeri & Steppan, 2018; Mason, 2016a). This may relate to the relative openness of the microhabitats used by *Antechinomys,* with the arid members of the genus preferring open stony plains over areas with higher vegetation cover (Dodd et al., 2025). This is consistent with patterns in rodent communities, where species with the largest bullae preferentially forage in open areas (Kotler, 1984).

Interspecific communication has been posited as playing a role in bullar hypertrophy in kangaroo rats (*Dipodomyinae*), which use low-frequency foot drumming to communicate (Randall, 1993, 1994); however, most taxa with large bullae show no evidence of low-frequency vocalizations. Given the near ubiquity of the pattern of aridity driving increases in bullar volume across taxonomically diverse groups of granivores (rodents), omnivores (rodents and bandicoots), and now insectivores (the present study), it seems likely that the shared selective pressure of predation is a key driver of this pattern. Prey detection and/or intraspecific communication may play a role in some species, and merits further study (Mason, 2016a).

Our results indicate that the orbital module tends to become narrower towards the equator in Sminthopsinae. This pattern is challenging to interpret as the orbital module is associated with multiple functions including sight, olfaction, cognition, and mastication (see Figure 1). This highlights the difficulty in interpreting skull shape evolution as differences in shape rarely pertain to a single aspect of an organism’s biology (Walker, 2007). We propose that the two most likely drivers of this pattern are thermoregulation and a tradeoff between eyesight and olfaction. Thermoregulation is a crucial challenge for small mammals due to their small surface area to volume ratio (Heldmaier, 1989), and many taxa possess behavioral or physiological adaptations for heat regulation (Degen, 1997; Terrien et al., 2011). The nasal cavity can play an important role in thermoregulation, with the complex nasal turbinals reabsorbing heat and moisture from exhaled air (Jackson & Schmidt-Nielsen, 1964; Schmidt-Nielsen et al., 1970). Our findings of a widening of the posterior nasal cavity in species further from the equator may be consistent with an increased demand for thermoregulatory capabilities relative to more tropical species (Boyles et al., 2013). This part of the nasal cavity is primarily occupied by olfactory, rather than respiratory turbinals (Macrini et al., 2023; Martinez et al., 2024), however turbinals can play a role in both processes and their functions are largely understudied (Martinez et al., 2024). Another factor to consider is the interaction between thermoregulation and the tradeoff between olfaction and vision (Chi et al., 2025; Nummela et al., 2013), as a wider posterior nasal cavity pushes the eyes apart, reducing the degree of binocular vision (Cartmill, 1972). As a result, it is possible that selection for improved thermoregulation at lower (more polar) latitudes pushes the vision-olfaction tradeoff towards olfaction due to the added thermoregulatory benefits of an increase in nasal volume. Additional work is needed on the roles of vision and olfaction in the sensory ecology of sminthopsins to determine the functional significance of this pattern.

The six-module system that best explained our dataset is largely similar to patterns of modularity found in the skulls of other mammal groups (Cheverud, 1982; Goswami, 2006; Heck et al., 2018; Randau et al., 2019; Sherratt & Kraatz, 2023). The main difference in our findings is that we found support for splitting the basicranium into two separate modules: a ‘bulla’ module and a ‘rest of the basicranium’ module. While not previously reported, this pattern is perhaps unsurprising given the functional differences between the sensory role of the bullae in hearing (Mason, 2016a) and the more structural role of the rest of the basicranium. This also highlights a key limitation in the modularity approach as it relies on comparing *a priori* grouping of landmarks into discrete modules, meaning that groupings not explicitly tested cannot be detected. We found the molar region (spanning the premolars and molars in our model) shows relatively little disparity, likely due to the dietary conservatism displayed amongst dasyurids. The basicranium module was the least integrated in our dataset, which is likely driven by the higher proportion of foramina-associated landmarks in our landmarking protocol for this module, which inherently covary less than other landmarks (Weisbecker et al., 2023).

## Conclusions

There is a growing understanding that the skull can play an important role in facilitating adaptation to the distinct sensory demands of arid environments, particularly in small mammals (Nummela et al., 2013). The openness of arid environments selects for improved long-distance sensory capabilities, which has manifested in improved low-frequency hearing in a range of taxa (Mason, 2016a). The Sminthopsinae show a clear pattern of increasing relative bulla size along an aridity gradient, leading to convergence in bulla shape amongst arid-adapted species across several independent biome transition events. Variation in skull shape can be highly complex. However, here we showed how the modularity framework can be used to identify the effects of module-specific selective pressures applied by the sensory (hearing) and physiological (thermoregulation) demands of arid environments. While hearing has likely played a key role in adaptation within the Sminthopsinae, additional work on inner ear morphology is needed to fully understand the complex role of sound and hearing in their sensory ecology and evolution.

