## Supplementary Material for "Auditory bullae enlarge along an aridity gradient in a diverse subfamily of small carnivorous mammals"

Supplementary methods

We expected the sminthopsin cranium to follow some variant on the six-module hypothesis that has been identified across many mammal groups (Kraatz & Sherratt, 2016; Marcus et al., 2000; Sherratt & Kraatz, 2023; Usui & Tokita, 2018). However, we wanted to comprehensively test a range of hypotheses to corroborate this. Due to the computational limitations on testing every possible combination of modules, our initial modularity testing focused on three well-established hypotheses: a two module tissue-of-origin hypothesis (Willmore et al., 2006; Zelditch, 1988), the 'Cheverud' six-module functional hypothesis (Cheverud, 1982; Moss, 1968; Willmore et al., 2006) (Willmore et al., 2006; Zelditch, 1988), and the 'Goswami' six-module functional hypothesis. Due to the proposed importance in the auditory bullae in adaptation to aridity, and aridity being a key driver of the radiation and hypothesized speciation of sminthopsins in the late Miocene (Archer, 1981; García-Navas et al., 2018), we also tested whether the bullae behaved as a separate module to the rest of the basicranium in sminthopsins by testing variants of the Cheverud and Goswami hypotheses in which the bullae were treated as a separate module. Due to the lack of landmarking along the premolar row in previous studies (or in the case of rodents, the absence of the premolar row) (Goswami, 2006; Sherratt & Kraatz, 2023), we also tested variants of the Goswami hypothesis in which premolar landmarks were assigned to either the rostrum or molar modules. This left seven initial hypotheses: tissue-of-origin, Cheverud (bullae as basicranium), Cheverud (bullae as separate), Goswami (bullae as basicranium + premolar as rostrum), Goswami (bullae as basicranium + premolar as molar), Goswami (bullae as separate + premolar as rostrum), Goswami (bullae as separate + premolar as molar). Details of landmark module assignments for each hypothesis are shown in Supplementary S3.

We tested our modularity hypotheses using EMMLi (Goswami & Finarelli, 2016), which uses a maximum likelihood approach to determine the best fitting modularity hypothesis and has the advantage of being able to test models with different numbers of modules. We generated vector congruence coefficient correlation matrices using *paleomorph::dotcorr* (Lucas & Goswami, 2017) to compare landmark covariance. This method treats each landmark as a point varying in 3D space rather than three points each varying in 1D space, which better reflects biological covariance patterns (Goswami & Finarelli, 2016).

Following an initial EMMLi analysis of the best fitting of our seven base modularity hypotheses, we inspected the within and between module correlations of the best-supported module to determine candidates for model simplification by merging modules (Bardua et al., 2019; Felice & Goswami, 2018). If a module pair had a plausible developmental or functional link and the between-module correlation was within 0.1 of either of the two within module values, following Felice and Goswami (2018), the modules were merged to generate an additional hypothesis. Following the generation of all alternate modularity hypotheses, we tested them using *EMMLiv2::subsampleEMMLi()* (Goswami et al., 2019; Goswami & Finarelli, 2016) with 100 replicates of 0.8 of the landmarks randomly subsampled for each replicate. We ran several variants of the analysis to ensure our choice of modularity hypothesis was robust. We analyzed the entire dataset as well as each of the four clades of sminthopsin separately. We performed both a single GPA analysis as well as module-specific GPAs, and using both allometry-corrected and allometry-non-corrected data.

Initial modularity hypothesis testing found strong support for the Goswami (bullae as separate + premolar as molar) hypothesis. Inspection of the within- and between- module variance of the best model sets revealed several module pairs and candidates for combining. However, ultimately the Goswami (bullae as separate + premolar as molar) hypothesis was still chosen as the best model in 45/50 parameter-dataset variants, including 10/10 using all genera pooled (Supplementary S4). More specifically, this hypothesis allowing for different magnitudes of within-module correlations was recovered in 37/45 analyses, so this was used as the best fit modularity hypothesis for further analyses.

Supplementary S1. List of crania analyzed from CT-scan data as part of this study. Species: *A.* = *Antechinomys*, *N.* = *Ningaui*, *S.* = *Sminthopsis*; ID: MAGNT = Museum and Art Gallery of the Northern Territory, AM = Australian Museum, SAMA = South Australian Museum and Art Gallery, QM = Queensland Museum, NMV = National Museum of Victoria, WAM = Western Australian Museum; Sex: U = unknown/inderterminable. Coordinates are presented to the decimal accuracy provided on the Atlas of Living Australia (ALA).

| Species | ID | Latitude | Longitude | Locality | Biome |
|---|---|---|---|---|---|
| *A. auritus* | MAGNT U2026 | -25.6 | 132.8167 | Erldunda Station WA | Hyper-arid |
| *A. auritus* | SAMA M7524 | -31.2 | 136.2333 | Wirraminna WA | Hyper-arid |
| *A. auritus* | WAM M23674 | -26.3666 | 117.3333 | Mileura Homestead WA | Hyper-arid |
| *A. auritus* | WAM M5775 | -26.1166 | 126.5833 | Elder Creek WA | Hyper-arid |
| *A. auritus* | WAM M5776 | -26.1166 | 126.5833 | Elder Creek WA | Hyper-arid |
| *A. auritus* | WAM M6355 | -25.8166 | 119 | New Springs Homestead WA | Hyper-arid |
| *A. auritus* | WAM M7738 | -26.0666 | 126.5833 | Warburton Range WA | Hyper-arid |
| *A. laniger* | AM M.8463 | -31.0833 | 146.4167 | El Trune Station NSW | Semi-arid |
| *A. laniger* | AM M.8464 | -31.0833 | 146.4167 | El Trune Station NSW | Semi-arid |
| *A. laniger* | AM M.8465 | -31.0833 | 146.4167 | El Trune Station NSW | Semi-arid |
| *A. laniger* | NMV C2537 | -22.63 | 136.8 | Tarlton Downs NSW | Semi-arid |
| *A. laniger* | QM JM1174 | -22.1 | 137.97 | Tobermorey NT | Semi-arid |
| *A. laniger* | QM JM1175 | -22.2833 | 137.97 | Tobermorey NT | Semi-arid |
| *A. laniger* | SAMA M1804 | -14.5333 | 135.25 | Roper River NT | Semi-arid |
| *A. spenceri* | NMV C7355 | -24.05 | 139.07 | Sandringham Station Qld | Hyper-arid |
| *A. spenceri* | QM JM1177 | -25.7 | 140.75 | Betoota Qld | Hyper-arid |
| *A. spenceri* | SAMA M22633 | -27.0175 | 138.8947 | Clifton Hills Station SA | Hyper-arid |
| *A. spenceri* | SAMA M22957 | -28.0167 | 138.6625 | Mungeranie Station SA | Hyper-arid |
| *A. spenceri* | SAMA M9485 | -27.0167 | 138.9 | Clifton Hills Station SA | Hyper-arid |
| *A. spenceri* | SAMA M9891 | -26.75 | 139.5667 | Clifton Hills Station SA | Hyper-arid |
| *A.* sp nov SW | WAM M20648 | -31.575 | 119.8667 | Toomey Hills WA | Mesic |
| *A.* sp nov SW | WAM M20649 | -31.5833 | 119.85 | Toomey Hills WA | Mesic |
| *A.* sp nov SW | WAM M2230 | -33.1 | 118.4667 | Lake Grace WA | Mesic |
| *A.* sp nov SW | WAM M2368 | -33.1 | 118.4667 | Lake Grace WA | Mesic |
| *A.* sp nov SW | WAM M53591 | -33.2572 | 121.0961 | Grass Patch WA | Mesic |
| *A. longicaudatus* | WAM M17424 | -24.5833 | 126.2833 | Clutterbuck Hills WA | Hyper-arid |
| *A. longicaudatus* | WAM M24100 | -25.0417 | 125 | Young Range WA | Hyper-arid |
| *A. longicaudatus* | WAM M41336 | -27.6733 | 119.4086 | Lake Mason Station WA | Hyper-arid |
| *A. longicaudatus* | WAM M46144 | NA | NA | WA | Hyper-arid |
| *A. longicaudatus* | WAM M56543 | -22.8322 | 120.4833 | Roy Hill Mine WA | Hyper-arid |
| *N. ridei* | WAM M20295 | -30.2917 | 119.7333 | Bungalbin Hill WA | Hyper-arid |
| *N. ridei* | WAM M20446 | -27.4 | 120.7 | Wanjarri Nature Reserve WA | Hyper-arid |
| *N. ridei* | WAM M20449 | -29.4833 | 119.6 | Mount Elvire WA | Hyper-arid |
| *N. ridei* | WAM M20451 | -29.4833 | 119.6 | Mount Elvire WA | Hyper-arid |
| *N. ridei* | WAM M20452 | -29.4833 | 119.6 | Mount Elvire WA | Hyper-arid |
| *N. ridei* | WAM M46747 | -21.6667 | 121.5833 | Plechtrachne WA | Hyper-arid |
| *N. timealeyi* | WAM M18142 | -22.3 | 113.8333 | Yardie Well WA | Hyper-arid |
| *N. timealeyi* | WAM M18675 | -22.8967 | 118.1272 | Hamersley River WA | Hyper-arid |
| *N. timealeyi* | WAM M19672 | -21.3 | 116.0278 | Mardie Homestead WA | Hyper-arid |
| *N. timealeyi* | WAM M23841 | -21.5833 | 117.0667 | Millstream WA | Hyper-arid |
| *N. timealeyi* | WAM M24590 | -21.25 | 116.1278 | Fortescue River WA | Hyper-arid |
| *N. timealeyi* | WAM M28211 | -21.6097 | 118.9622 | Woodstock WA | Hyper-arid |

| | | | | | |
|---|---|---|---|---|---|
| *N. timealeyi* | WAM M45033 | -22.2028 | 114.0403 | Cape Range WA | Hyper-arid |
| *N. yvonneae* | WAM M23391 | -29.8583 | 123.3778 | Queen Victoria Springs WA | Semi-arid |
| *N. yvonneae* | WAM M40561 | -30.75 | 121.4667 | Kalgoorlie WA | Semi-arid |
| *N. yvonneae* | WAM M40686 | -30.75 | 121.4667 | Kalgoorlie WA | Semi-arid |
| *N. yvonneae* | WAM M40698 | -30.3 | 119.7333 | Manjimup WA | Semi-arid |
| *N. yvonneae* | WAM M47074 | -30.3 | 119.7333 | Bungalbin Hill WA | Semi-arid |
| *N. yvonneae* | WAM M49423 | -29.9456 | 121.1208 | Goongarrie Station WA | Semi-arid |
| *S. archeri* | QM JM10027 | -12.6964 | 143.3331 | Ham Hill WA | Mesic |
| *S. archeri* | QM JM15794 | -19.5511 | 144.06 | Blackbraes NP Qld | Mesic |
| *S. butleri* | SAMA M4072 | -11.5833 | 131.1667 | Melville Island NT | Mesic |
| *S. butleri* | WAM M7158 | -14.3 | 126.7 | Kalumburu WA | Mesic |
| *S. dolichura* | WAM M13141 | -31.4 | 117.6583 | Yorkrakine WA | Semi-arid |
| *S. dolichura* | WAM M26194 | -28.5667 | 119.0167 | Yuinmery Station WA | Semi-arid |
| *S. dolichura* | WAM M26733 | -29.9667 | 123.7667 | Yuinmery Station WA | Semi-arid |
| *S. dolichura* | WAM M29940 | -30.25 | 115.0167 | Mount Lereuer NP WA | Semi-arid |
| *S. dolichura* | WAM M40544 | -33.95 | 115.0667 | Margaret River WA | Semi-arid |
| *S. dolichura* | WAM M47534 | NA | NA | Janderup WA | Semi-arid |
| *S. fuliginosa* | WAM M23821 | -34.4292 | 115.7417 | Black Point WA | Mesic |
| *S. fuliginosa* | WAM M26801 | -34.3667 | 118.25 | Stirling Range NP WA | Mesic |
| *S. fuliginosa* | WAM M29128 | -35.1167 | 117.9167 | Torndirrup NP WA | Mesic |
| *S. fuliginosa* | WAM M30911 | -33.9667 | 119.3333 | Fitzgerald River NP WA | Mesic |
| *S. fuliginosa* | WAM M47598 | -34.3333 | 116.25 | Manjimup WA | Mesic |
| *S. fuliginosa* | WAM M52167 | -33.8222 | 123.1972 | Cape Arid NP WA | Mesic |
| *S. gilberti* | WAM M23137 | -31.9 | 116.6333 | Wambin Reserve WA | Mesic |
| *S. gilberti* | WAM M23664 | -32.725 | 116.35 | Boddington Gold Mine WA | Mesic |
| *S. gilberti* | WAM M26116 | -32.15 | 116.1333 | Canning River WA | Mesic |
| *S. gilberti* | WAM M27260 | -32.55 | 117.3333 | Tutanning Reserve WA | Mesic |
| *S. gilberti* | WAM M41897 | NA | NA | Dobadelly Nature Reserve WA | Mesic |
| *S. gilberti* | WAM M44035 | -34.3028 | 116.2294 | Manjimup WA | Mesic |
| *S. granulipes* | WAM M17501 | -32.4042 | 119.6861 | Lake Cronin WA | Mesic |
| *S. granulipes* | WAM M17649 | -32.3875 | 119.6861 | Lake Cronin WA | Mesic |
| *S. granulipes* | WAM M53203 | -33.0814 | 119.9978 | Lake King WA | Mesic |
| *S. granulipes* | WAM M53589 | -33.0419 | 119.9914 | Lake King WA | Mesic |
| *S. granulipes* | WAM M55479 | -33.3475 | 120.8717 | Grass Patch WA | Mesic |
| *S. hirtipes* | WAM M15261 | -28.34 | 123.1083 | White Cliffs Homestead WA | Hyper-arid |
| *S. hirtipes* | WAM M1577 | -26.525 | 126.025 | Winduldurra Rockhole WA | Hyper-arid |
| *S. hirtipes* | WAM M23353 | -29.925 | 123.5278 | Queen Victoria Springs WA | Hyper-arid |
| *S. hirtipes* | WAM M23356 | -29.8306 | 123.35 | Queen Victoria Springs WA | Hyper-arid |
| *S. hirtipes* | WAM M25953 | -28.3 | 125.8167 | Neale Junction WA | Hyper-arid |
| *S. hirtipes* | WAM M26948 | -30.4333 | 123.5667 | Queen Victoria Springs WA | Hyper-arid |
| *S. leucopus* | AM M.12954 | -36.983 | 149.55 | Bombala NSW | Mesic |
| *S. leucopus* | AM M.13271 | -36.55 | 149.866 | Mumbulla State Forest NSW | Mesic |
| *S. leucopus* | AM M.13272 | -36.55 | 149.866 | Mumbulla State Forest NSW | Mesic |
| *S. leucopus* | AM M.17302 | -36.833 | 149.583 | Coolangubra State Forest NSW | Mesic |
| *S. leucopus* | AM M.4343 | -40.533 | 148.166 | Clarke Island Tas | Mesic |
| *S. leucopus* | AM M.7425 | -38.35 | 141.6 | South Portland Vic | Mesic |
| *S. murina* | AM M.35820 | NA | NA | NA | Mesic |
| *S. murina* | AM M.6639 | -33.516 | 150.766 | East Kurrajong NSW | Mesic |

| | | | | | |
|---|---|---|---|---|---|
| *S. murina* | AM M.7402 | -29.6 | 150.65 | Warialda NSW | Mesic |
| *S. murina* | AM M.9835 | -28.833 | 149 | Mungindi NSW | Mesic |
| *S. murina* | WAM M21326 | -32.3833 | 152.4833 | Smith's Lake NSW | Mesic |
| *S. murina* | WAM M35633 | NA | NA | Penda State Forrest, SA | Mesic |
| *S. ooldea* | WAM M18243 | -23.1847 | 118.7919 | Mount Meharry WA | Hyper-arid |
| *S. ooldea* | WAM M23660 | -28.2333 | 125.8667 | Neale Junction WA | Hyper-arid |
| *S. ooldea* | WAM M43222 | -22.6022 | 118.5208 | Mount Windell WA | Hyper-arid |
| *S. ooldea* | WAM M49459 | -26.2261 | 121.5189 | Lorna Glen Station WA | Hyper-arid |
| *S. ooldea* | WAM M52173 | -23.0978 | 118.8386 | West Angelas Mine WA | Hyper-arid |
| *S. ooldea* | WAM M62093 | -27.2386 | 120.7328 | Lake Way WA | Hyper-arid |
| *S. ooldea* | WAM M8077 | -30.15 | 132.2167 | Maralinga SA | Hyper-arid |
| *S. psammophila* | WAM M23226 | -29.9 | 123.5 | Queen Victoria Springs WA | Semi-arid |
| *S. psammophila* | WAM M23227 | -29.9 | 123.6 | Queen Victoria Springs WA | Semi-arid |
| *S. psammophila* | WAM M23228 | -29.9 | 123.6 | Queen Victoria Springs WA | Semi-arid |
| *S. psammophila* | WAM M23229 | -30.1 | 123.8 | Queen Victoria Springs WA | Semi-arid |
| *S. psammophila* | WAM M23230 | -30 | 123.6 | Queen Victoria Springs WA | Semi-arid |
| *S. youngsoni* | WAM M22621 | -22.45 | 123.9333 | Great Sandy Desert WA | Hyper-arid |
| *S. youngsoni* | WAM M25604 | -23.1 | 123.3667 | Canning Stock Route WA | Hyper-arid |
| *S. youngsoni* | WAM M25774 | -22.3294 | 122.0692 | Rudall River NP WA | Hyper-arid |
| *S. youngsoni* | WAM M25777 | -22.3431 | 122.0683 | Rudall River NP WA | Hyper-arid |
| *S. youngsoni* | WAM M25780 | -22.3247 | 122.0756 | Rudall River NP WA | Hyper-arid |
| *S. youngsoni* | WAM M29310 | -22.3042 | 122.0447 | Kintyre WA | Hyper-arid |
| *S. youngsoni* | WAM M48509 | -25.7108 | 114.5997 | Meedo Homestead WA | Hyper-arid |
| *S. youngsoni* | WAM M48785 | NA | NA | Great Victoria Desert WA | Hyper-arid |
| *S. bindi* | AM M36287 | -13.825 | 131.833 | Pine Creek NT | Mesic |
| *S. crassicaudata* | WAM M20411 | -30.55 | 121.2333 | Kalgoorlie WA | Semi-arid |
| *S. crassicaudata* | WAM M28294 | -31.8167 | 128.2167 | Nullarbor Plain WA | Semi-arid |
| *S. crassicaudata* | WAM M28334 | -31.8167 | 128.2167 | Nullarbor Plain WA | Semi-arid |
| *S. crassicaudata* | WAM M4491 | -33.6833 | 136.9167 | Eyre Peninsula SA | Semi-arid |
| *S. crassicaudata* | WAM M4504 | -29.0167 | 138.4667 | Dulkaninna Homestead SA | Semi-arid |
| *S. crassicaudata* | WAM M7746 | -31 | 124.6833 | Maretha Siding WA | Semi-arid |
| *S. crassicaudata* | WAM M7896 | -27.7528 | 114.3583 | Kalbarri NP WA | Semi-arid |
| *S. crassicaudata* | WAM M8084 | -37.2833 | 142.9333 | Ararat VIC | Semi-arid |
| *S. douglasi* | QM JM18109 | -19.4667 | 141.2 | Lyrian Station Qld | Semi-arid |
| *S. douglasi* | QM JM18110 | -19.4667 | 141.2 | Lyrian Station Qld | Semi-arid |
| *S. douglasi* | QM JM18111 | -19.4667 | 141.2 | Lyrian Station Qld | Semi-arid |
| *S. douglasi* | QM JM18113 | -19.4667 | 141.2 | Lyrian Station Qld | Semi-arid |
| *S. douglasi* | QM JM18115 | -21.4333 | 142.1167 | Crendon Station Qld | Semi-arid |
| *S. douglasi* | QM JM18600 | -20.9 | 142.1333 | Proa via Nelia | Semi-arid |
| *S. froggatti* | WAM M18832 | -16.4736 | 128.5583 | Lissadell Station WA | Mesic |
| *S. froggatti* | WAM M19274 | -16.6389 | 128.45 | Lissadell Station WA | Mesic |
| *S. froggatti* | WAM M24390 | -16.3833 | 126.2667 | Mount Elizabeth WA | Mesic |
| *S. macroura N* | QM J11133 | -25.6167 | 151.6167 | Gayndah Qld | Semi-arid |
| *S. macroura N* | QM J8179 | -24.4667 | 150.1333 | Banana Qld | Semi-arid |
| *S. macroura N* | QM JM16573 | -20.8333 | 144.5 | Charters Towers Qld | Semi-arid |
| *S. macroura N* | QM JM19215 | -27.1 | 151.54 | Moola Qld | Semi-arid |
| *S. macroura N* | QM JM986 | -26.6333 | 145.0167 | Cheepie Qld | Semi-arid |
| *S. macroura N* | WAM M4511 | -20.5667 | 147.0833 | Collinville Qld | Semi-arid |

| | | | | | |
|---|---|---|---|---|---|
| *S. macroura S* | SAMA M11615 | -29.1583 | 134.3333 | Mabel Creek Homestead SA | Semi-arid |
| *S. macroura S* | SAMA M12368 | -32.15 | 135.1917 | Peeweena Dam SA | Semi-arid |
| *S. macroura S* | SAMA M14911 | -31.2 | 136.82 | Woomera SA | Semi-arid |
| *S. macroura S* | SAMA M18506 | -29.5138 | 136.1563 | Johnson Bore WA | Semi-arid |
| *S. macroura S* | SAMA M7931 | -28.3 | 136.5 | Johnson WA | Semi-arid |
| *S. macroura S* | SAMA M7932 | -28.3 | 136.5 | Johnson WA | Semi-arid |
| *S. macroura W* | WAM M5698 | -18.5278 | 125.9056 | Gogo Station WAM | Hyper-arid |
| *S. macroura W* | WAM M19817 | -21.0014 | 117.1222 | Roebourne WA | Hyper-arid |
| *S. macroura W* | WAM M22692 | -22.45 | 123.9 | Great Sandy Desert WA | Hyper-arid |
| *S. macroura W* | WAM M47690 | -22.8086 | 117.7789 | Tom Price WA | Hyper-arid |
| *S. macroura W* | WAM M47842 | -22.8178 | 117.7639 | Tom Price WA | Hyper-arid |
| *S. macroura W* | WAM M5086 | -22.7167 | 119.5667 | Coondiner Pool WA | Hyper-arid |
| *S. macroura W* | WAM M6075 | -27.9972 | 122.8931 | Cosmo Newberry WA | Hyper-arid |
| *S. macroura W* | WAM M16758 | -18.9083 | 127.7167 | Wolf Creek WA | Hyper-arid |
| *S. nitela* | WAM M15785 | -14.6694 | 125.7389 | Mitchell Plateau WA | Mesic |
| *S. nitela* | WAM M15786 | -14.6722 | 125.7361 | Mitchell Plateau WA | Mesic |
| *S. nitela* | WAM M21514 | -14.8792 | 125.8083 | Mitchell Plateau WA | Mesic |
| *S. nitela* | WAM M21515 | -14.8222 | 125.8417 | Mitchell Plateau WA | Mesic |
| *S. nitela* | WAM M21996 | -14.8167 | 125.8403 | Mitchell Plateau WA | Mesic |
| *S. nitela* | WAM M4056 | -12.3167 | 133.05 | Gunbalanya NT | Mesic |
| *S. stalkeri* | SAMA M8339 | -22.25 | 131.8 | Yuendumu NT | Semi-arid |
| *S. stalkeri* | SAMA M8292 | -17.9333 | 138.8167 | Doomadgee Qld | Semi-arid |
| *S. stalkeri* | SAMA M8293 | -17.9333 | 138.8167 | Doomadgee Qld | Semi-arid |
| *S. stalkeri* | SAMA M8295 | -17.9333 | 138.8167 | Doomadgee Qld | Semi-arid |
| *S. stalkeri* | SAMA M8296 | -17.9333 | 138.8167 | Doomadgee Qld | Semi-arid |
| *S. stalkeri* | SAMA M8326 | -17.9333 | 138.8167 | Doomadgee Qld | Semi-arid |
| *S. virginiae* | QM JM4017 | -11.983 | 141.8691 | Mapoon Plains Qld | Mesic |
| *S. virginiae* | QM JM4018 | -12.5402 | 141.7095 | Pine River Bay Qld | Mesic |
| *S. virginiae* | QM JM4019 | -12.7517 | 142.3111 | York Downs Qld | Mesic |
| *S. virginiae* | QM JM689 | -15.75 | 145.2833 | Home Rule Qld | Mesic |

Supplementary S2. A list of the landmarks used in this study. IDs match Figure 1A.

| Landmark | Description |
|---|---|
| LM001_C | Midline point on the premaxilla between the upper incisors |
| LM003_R | Frontal-maxillary-nasal intersection |
| LM004_C | Nasal-frontal suture midline |
| LM005_L | Frontal-maxillary-nasal intersection |
| LM006_R | Most antero-distal point in upper first molar neck |
| LM007_R | Most anterolateral point in lacrimal ridge at midpoint of lacrimal foramen |
| LM008_R | Posterior tip of lacrimal |
| LM009_L | Most antero-distal point in upper first molar neck |
| LM010_L | Most anterolateral point in lacrimal ridge at midpoint of lacrimal foramen |
| LM011_L | Posterior tip of lacrimal |
| LM012_R | Most posterior point in the jugal-maxillary suture on zygomatic |
| LM013_L | Most posterior point in the jugal-maxillary suture on zygomatic |
| LM014_C | Frontal-parietal suture midline |
| LM015_R | Supraoccipital-petrosal-squamosal intersection on nuchal crest |
| LM016_C | External occipital protuberance nuchal crest midline |
| LM017_L | Supraoccipital-petrosal-squamosal intersection on nuchal crest |
| LM018_C | Dorsal lip of foramen magnum midline |
| LM019_R | Maximum concavity in distal of foramen magnum |
| LM020_C | Ventral lip of foramen magnum midline |
| LM021_L | Maximum concavity in distal of foramen magnum |
| LM022_R | Paracondylar process of the exoccipital ventral |
| LM023_R | Point of contact between petrosal and ventrolateral edge of alisphenoid |
| LM024_R | Most ventral point on tympanic bulla |
| LM025_L | Paracondylar process of the exoccipital ventral |
| LM026_L | Point of contact between petrosal and ventrolateral edge of alisphenoid |
| LM027_L | Most ventral point on tympanic bulla |
| LM028_C | Palatine posterior midline |
| LM029_R | Major palatine foramen posterior |
| LM030_L | Major palatine foramen posterior |
| LM031_R | Incisive foramen posterior |
| LM032_L | Incisive foramen posterior |
| LM033_R | Most posterolateral point in jugal-squamosal suture |
| LM034_R | Anterior-most point of lateral tympanic bulla edge |
| LM035_R | Postglenoid process ventral |
| LM036_R | Suprameatal foramen posterior |
| LM037_L | Most posterolateral point in jugal-squamosal suture |
| LM038_L | Anterior-most point of lateral tympanic bulla edge |
| LM039_L | Postglenoid process ventral |
| LM040_L | Suprameatal foramen posterior |
| LM041_R | Frontal-maxillary-lacrimal intersection |
| LM042_L | Frontal-maxillary-lacrimal intersection |
| LM047_R | Maximum concavity on anterior edge of premaxillary |
| LM048_L | Maximum concavity on anterior edge of premaxillary |
| LM049_R | Premaxillary-maxillary suture anterior lateral end |
| LM050_L | Premaxillary-maxillary suture anterior lateral end |

| | |
|---|---|
| LM051_R | Premaxillary-maxillary suture posterior |
| LM052_L | Premaxillary-maxillary suture posterior |
| LM053_R | Most posterio-distal point in upper canine neck |
| LM054_L | Most posterio-distal point in upper canine neck |
| LM055_R | Most ventral point of infraorbital foramen |
| LM056_R | Most dorsal point of infraorbital foramen |
| LM057_L | Most ventral point of infraorbital foramen |
| LM058_L | Most dorsal point of infraorbital foramen |
| LM059_R | Maxillary-lacrimal-jugal intersection |
| LM060_L | Maxillary-lacrimal-jugal intersection |
| LM061_R | Most posterior point in fourth molar neck |
| LM062_L | Most posterior point in fourth molar neck |
| LM063_R | Most posteromedial point in the jugal-squamosal suture |
| LM064_L | Most posteromedial point in the jugal-squamosal suture |
| LM065_R | Suprameatal foramen anterior |
| LM066_L | Suprameatal foramen anterior |
| LM067_R | Nuchal crest-posttypanic process intersection |
| LM068_L | Nuchal crest-posttypanic process intersection |
| LM071_R | Most medial point in hypoglossal foramen |
| LM072_L | Most medial point in hypoglossal foramen |
| LM073_R | Most anterior point in jugular foramen |
| LM074_L | Most anterior point in jugular foramen |
| LM075_R | Most anterior point of the basioccipital |
| LM076_L | Most anterior point of the basioccipital |
| LM077_R | Most anterior point in carotid canal |
| LM078_L | Most anterior point in carotid canal |
| LM079_R | Major palatine foramen anterior |
| LM080_L | Major palatine foramen anterior |
| LM081_R | Incisive foramen anterior |
| LM082_L | Incisive foramen anterior |
| LM083_R | Most anterior point of frontal-parietal-squamosal suture |
| LM084_L | Most anterior point of frontal-parietal-squamosal suture |
| LM085_R | Lateral-most point of tympanic bulla |
| LM086_L | Lateral-most point of tympanic bulla |
| LM087_R | Most dorsal point on tympanic bulla |
| LM088_L | Most dorsal point on tympanic bulla |
| LM089_R | Anterior-most point of medial tympanic bulla edge |
| LM090_L | Anterior-most point of medial tympanic bulla edge |
| LM091_R | Most dorsoanterior point of foramen ovale |
| LM092_L | Most dorsoanterior point of foramen ovale |
| LM093_R | Transverse canal posterior |
| LM094_L | Transverse canal posterior |
| LM095_R | Transverse canal anterior |
| LM096_L | Transverse canal anterior |
| LM097_R | Foramen rotundum lateral |
| LM098_L | Foramen rotundum lateral |
| LM099_R | anterior-most point of interior of tympanic bulla |
| LM100_L | anterior-most point of interior of tympanic bulla |

| | |
|---|---|
| LM101_R | lateral-most point of post-orbital foramen |
| LM102_L | lateral-most point of post-orbital foramen |
| C01_C | LM003 to LM004. midline. nasal suture (7) |
| C02_R | LM043 to LM051. nasal-premaxilla suture (5) |
| C03_L | LM044 to LM052. nasal-premaxilla suture (5) |
| C04_R | LM051 to LM003. nasal-maxilla suture (5) |
| C05_L | LM052 to LM005. nasal-maxilla suture (5) |
| C06_R | LM004 to LM003. nasal-frontal suture (4) |
| C07_L | LM004 to LM005. nasal-frontal suture (4) |
| C08_R | LM049 to LM051. premaxilla-maxilla suture (7) |
| C09_L | LM050 to LM052. premaxilla-maxilla suture (7) |
| C10_R | LM003 to LM041. frontal-maxilla suture (7) |
| C11_L | LM005 to LM042. frontal-maxilla suture (7) |
| C12_R | LM008 to LM007. anterior lacrimal edge (7) |
| C13_L | LM011 to LM010. anterior lacrimal edge (7) |
| C14_C | LM004 to LM014. midline. frontal suture (11) |
| C15_R | LM014 to LM083. frontal-parietal suture (9) |
| C16_L | LM014 to LM084. frontal-parietal suture (9) |
| C17_C | LM014 to LM016. midline. parietal suture (9) |
| C18_R | LM016 to LM015. parietal-occipital suture. on nuchal crest (7) |
| C19_L | LM016 to LM017. parietal-occipital suture. on nuchal crest (7) |
| C20_R | LM015 to LM067. parietal-petrosal suture. on nuchal crest (7) |
| C21_L | LM017 to LM068. parietal-petrosal suture. on nuchal crest (7) |
| C22_R | LM018 to LM019. foramen magnum dorsolateral edge (4) |
| C23_L | LM018 to LM021. foramen magnum dorsolateral edge (4) |
| C24_R | LM020 to LM019. foramen magnum ventrolateral edge (5) |
| C25_L | LM020 to LM021. foramen magnum ventrolateral edge (5) |
| C26_R | LM073 to LM075. basioccipital-petrosal suture (7) |
| C27_L | LM074 to LM076. basioccipital-petrosal suture (7) |
| C28_R | LM075 to LM023. petrosal-alisphenoid suture. on bulla (7) |
| C29_L | LM076 to LM026. petrosal-alisphenoid suture. on bulla (7) |
| C30_R | LM085 to LM099. anterior bulla-alisphenoid crease (9) |
| C31_L | LM086 to LM100. anterior bulla-alisphenoid crease (9) |
| C32_R | LM001 to LM049. premaxilla ventral edge (7) |
| C33_L | LM001 to LM050. premaxilla ventral edge (7) |
| C34_R | LM053 to LM006. maxilla ventral edge at premolars (7) |
| C35_L | LM054 to LM009. maxilla ventral edge at premolars (7) |
| C36_R | LM006 to LM061. maxilla ventral edge at molars (9) |
| C37_L | LM009 to LM062. maxilla ventral edge at molars (9) |

Supplementary S3. Showing each cranial modularity hypothesis tested in this study and the which landmarks were assigned to each module. Abbreviations are as follows: p = point landmark, c = curve (semi)landmark, nc = developmentally derived from the neural crest, me = developmentally derived from the mesoderm, f = face, v = vault, b = basicranium (including bullae), ba = basicranium (excluding bullae), bu = bullae only, t = temporal, r = rostrum, o = orbital, m = molar, or = orbital-rostrum combined module, vba = vault-basicranium (excluding bullae) combined module.

| name | Landmark type | Developmental | Cherverud | | Goswami | | | | | | | | | | |
|---|---|---|---|---|---|---|---|---|---|---|---|---|---|---|---|
| | | tissue-of-origin | bullae as basicranium | bullae as separate | bullae as basicranium+premolar as rostrum | bullae as basicranium+ premolar as molar | | bullae as separate+premolar as rostrum | | | | bullae as separate+premolar as molar | | | |
| | | base model | base model | base model | base model | base model | orbital and rostrum combined | base model | orbital and rostrum combined | vault and basicranium combined | vault and basicranium combined+orbital and rostrum combined | base model | orbital and rostrum combined | vault and basicranium combined | vault and basicranium combined+orbital and rostrum combined |
| LM001_C | p | nc | f | f | r | r | or | r | or | r | or | r | or | r | or |
| LM003_R | p | nc | f | f | o | o | or | o | or | o | or | o | or | o | or |
| LM004_C | p | nc | f | f | o | o | or | o | or | o | or | o | or | o | or |
| LM005_L | p | nc | f | f | o | o | or | o | or | o | or | o | or | o | or |
| LM006_R | p | nc | f | f | m | m | m | m | m | m | m | m | m | m | m |
| LM007_R | p | nc | t | t | o | o | or | o | or | o | or | o | or | o | or |
| LM008_R | p | m | t | t | o | o | or | o | or | o | or | o | or | o | or |
| LM009_L | p | nc | f | f | m | m | m | m | m | m | m | m | m | m | m |
| LM010_L | P | nc | t | t | o | o | or | o | or | o | or | o | or | o | or |
| LM011_L | p | nc | t | t | o | o | or | o | or | o | or | o | or | o | or |
| LM012_R | p | nc | t | t | NA | NA | NA | NA | NA | NA | NA | NA | NA | NA | NA |
| LM013_L | p | nc | t | t | NA | NA | NA | NA | NA | NA | NA | NA | NA | NA | NA |
| LM014_C | p | me | v | v | v | v | v | v | v | vba | vba | v | v | vba | vba |
| LM015_R | p | me | v | v | v | v | v | v | v | vba | vba | v | v | vba | vba |
| LM016_C | p | me | v | v | v | v | v | v | v | vba | vba | v | v | vba | vba |
| LM017_L | p | me | v | v | v | v | v | v | v | vba | vba | v | v | vba | vba |
| LM018_C | p | me | b | ba | b | b | b | ba | ba | vba | vba | ba | ba | vba | vba |
| LM019_R | p | me | b | ba | b | b | b | ba | ba | vba | vba | ba | ba | vba | vba |
| LM020_C | p | me | b | ba | b | b | b | ba | ba | vba | vba | ba | ba | vba | vba |
| LM021_L | p | me | b | ba | b | b | b | ba | ba | vba | vba | ba | ba | vba | vba |

| | | | | | | | | | | | | | | | |
|---|---|---|---|---|---|---|---|---|---|---|---|---|---|---|---|
| LM022_R | p | me | b | ba | b | b | b | ba | ba | vba | vba | ba | ba | vba | vba |
| LM023_R | p | me | b | bu | b | b | b | bu | bu | bu | bu | bu | bu | bu | bu |
| LM024_R | P | me | b | bu | b | b | b | bu | bu | bu | bu | bu | bu | bu | bu |
| LM025_L | p | me | b | ba | b | b | b | ba | ba | vba | vba | ba | ba | vba | vba |
| LM026_L | p | me | b | bu | b | b | b | bu | bu | bu | bu | bu | bu | bu | bu |
| LM027_L | p | me | b | bu | b | b | b | bu | bu | bu | bu | bu | bu | bu | bu |
| LM028_C | p | nc | f | f | m | m | m | m | m | m | m | m | m | m | m |
| LM029_R | p | nc | f | f | m | m | m | m | m | m | m | m | m | m | m |
| LM030_L | p | nc | f | f | m | m | m | m | m | m | m | m | m | m | m |
| LM031_R | p | nc | f | f | r | r | or | r | or | r | or | r | or | r | or |
| LM032_L | p | nc | f | f | r | r | or | r | or | r | or | r | or | r | or |
| LM033_R | p | nc | t | t | NA | NA | NA | NA | NA | NA | NA | NA | NA | NA | NA |
| LM034_R | p | me | b | bu | b | b | b | bu | bu | bu | bu | bu | bu | bu | bu |
| LM035_R | p | me | b | ba | b | b | b | ba | ba | vba | vba | ba | ba | vba | vba |
| LM036_R | p | nc | v | v | v | v | v | v | v | vba | vba | v | v | vba | vba |
| LM037_L | p | nc | t | t | NA | NA | NA | NA | NA | NA | NA | NA | NA | NA | NA |
| LM038_L | P | me | b | bu | b | b | b | bu | bu | bu | bu | bu | bu | bu | bu |
| LM039_L | p | me | b | ba | b | b | b | ba | ba | vba | vba | ba | ba | vba | vba |
| LM040_L | p | nc | v | v | v | v | v | v | v | vba | vba | v | v | vba | vba |
| LM041_R | p | me | v | v | o | o | or | o | or | o | or | o | or | o | or |
| LM042_L | p | me | v | v | o | o | or | o | or | o | or | o | or | o | or |
| LM047_R | p | nc | f | f | r | r | or | r | or | r | or | r | or | r | or |
| LM048_L | p | nc | f | f | r | r | or | r | or | r | or | r | or | r | or |
| LM049_R | p | nc | f | f | r | r | or | r | or | r | or | r | or | r | or |
| LM050_L | p | nc | f | f | r | r | or | r | or | r | or | r | or | r | or |
| LM051_R | p | nc | f | f | r | r | or | r | or | r | or | r | or | r | or |
| LM052_L | P | nc | f | f | r | r | or | r | or | r | or | r | or | r | or |
| LM053_R | p | nc | f | f | r | r | or | r | or | r | or | r | or | r | or |
| LM054_L | p | nc | f | f | r | r | or | r | or | r | or | r | or | r | or |
| LM055_R | p | nc | f | f | r | m | m | r | or | r | or | m | m | m | m |
| LM056_R | p | nc | f | f | r | m | m | r | or | r | or | m | m | m | m |

| | | | | | | | | | | | | | | | |
|---|---|---|---|---|---|---|---|---|---|---|---|---|---|---|---|
| LM057_L | p | nc | f | f | r | m | m | r | or | r | or | m | m | m | m |
| LM058_L | p | nc | f | f | r | m | m | r | or | r | or | m | m | m | m |
| LM059_R | p | nc | t | t | o | o | or | o | or | o | or | o | or | o | or |
| LM060_L | p | nc | t | t | o | o | or | o | or | o | or | o | or | o | or |
| LM061_R | p | nc | f | f | m | m | m | m | m | m | m | m | m | m | m |
| LM062_L | p | nc | f | f | m | m | m | m | m | m | m | m | m | m | m |
| LM063_R | p | nc | t | t | NA | NA | NA | NA | NA | NA | NA | NA | NA | NA | NA |
| LM064_L | p | nc | t | t | NA | NA | NA | NA | NA | NA | NA | NA | NA | NA | NA |
| LM065_R | p | nc | v | v | v | v | v | v | v | vba | vba | v | v | vba | vba |
| LM066_L | p | nc | v | v | v | v | v | v | v | vba | vba | v | v | vba | vba |
| LM067_R | p | me | v | v | v | v | v | v | v | vba | vba | v | v | vba | vba |
| LM068_L | p | me | v | v | v | v | v | v | v | vba | vba | v | v | vba | vba |
| LM071_R | p | me | b | ba | b | b | b | ba | ba | vba | vba | ba | ba | vba | vba |
| LM072_L | p | me | b | ba | b | b | b | ba | ba | vba | vba | ba | ba | vba | vba |
| LM073_R | p | me | b | ba | b | b | b | ba | ba | vba | vba | ba | ba | vba | vba |
| LM074_L | p | me | b | ba | b | b | b | ba | ba | vba | vba | ba | ba | vba | vba |
| LM075_R | p | me | b | bu | b | b | b | bu | bu | bu | bu | bu | bu | bu | bu |
| LM076_L | p | me | b | bu | b | b | b | bu | bu | bu | bu | bu | bu | bu | bu |
| LM077_R | p | me | b | ba | b | b | b | ba | ba | vba | vba | ba | ba | vba | vba |
| LM078_L | p | me | b | ba | b | b | b | ba | ba | vba | vba | ba | ba | vba | vba |
| LM079_R | p | nc | f | f | m | m | m | m | m | m | m | m | m | m | m |
| LM080_L | p | nc | f | f | m | m | m | m | m | m | m | m | m | m | m |
| LM081_R | p | nc | f | f | r | r | or | r | or | r | or | r | or | r | or |
| LM082_L | p | nc | f | f | r | r | or | r | or | r | or | r | or | r | or |
| LM083_R | p | me | t | t | v | v | v | v | v | vba | vba | v | v | vba | vba |
| LM084_L | P | me | t | t | v | v | v | v | v | vba | vba | v | v | vba | vba |
| LM085_R | p | me | b | bu | b | b | b | bu | bu | bu | bu | bu | bu | bu | bu |
| LM086_L | p | me | b | bu | b | b | b | bu | bu | bu | bu | bu | bu | bu | bu |
| LM087_R | p | me | b | bu | b | b | b | bu | bu | bu | bu | bu | bu | bu | bu |
| LM088_L | p | me | b | bu | b | b | b | bu | bu | bu | bu | bu | bu | bu | bu |
| LM089_R | p | me | b | bu | b | b | b | bu | bu | bu | bu | bu | bu | bu | bu |

| | | | | | | | | | | | | | | | |
|---|---|---|---|---|---|---|---|---|---|---|---|---|---|---|---|
| LM090_L | p | me | b | bu | b | b | b | bu | bu | bu | bu | bu | bu | bu | bu |
| LM091_R | p | me | b | ba | b | b | b | ba | ba | vba | vba | ba | ba | vba | vba |
| LM092_L | p | me | b | ba | b | b | b | ba | ba | vba | vba | ba | ba | vba | vba |
| LM093_R | p | me | b | ba | b | b | b | ba | ba | vba | vba | ba | ba | vba | vba |
| LM094_L | p | me | b | ba | b | b | b | ba | ba | vba | vba | ba | ba | vba | vba |
| LM095_R | p | me | b | ba | b | b | b | ba | ba | vba | vba | ba | ba | vba | vba |
| LM096_L | p | me | b | ba | b | b | b | ba | ba | vba | vba | ba | ba | vba | vba |
| LM097_R | p | me | b | ba | b | b | b | ba | ba | vba | vba | ba | ba | vba | vba |
| LM098_L | P | me | b | ba | b | b | b | ba | ba | vba | vba | ba | ba | vba | vba |
| LM099_R | p | me | b | bu | b | b | b | bu | bu | bu | bu | bu | bu | bu | bu |
| LM100_L | p | me | b | bu | b | b | b | bu | bu | bu | bu | bu | bu | bu | bu |
| LM101_R | p | me | v | v | o | o | or | o | or | o | or | o | or | o | or |
| LM102_L | p | me | v | v | o | o | or | o | or | o | or | o | or | o | or |
| C01_C | c | nc | f | f | r | r | or | r | or | r | or | r | or | r | or |
| C02_R | c | nc | f | f | r | r | or | r | or | r | or | r | or | r | or |
| C03_L | c | nc | f | f | r | r | or | r | or | r | or | r | or | r | or |
| C04_R | c | nc | f | f | r | r | or | r | or | r | or | r | or | r | or |
| C05_L | c | nc | f | f | r | r | or | r | or | r | or | r | or | r | or |
| C06_R | c | nc | f | f | o | o | or | o | or | o | or | o | or | o | or |
| C07_L | c | nc | f | f | o | o | or | o | or | o | or | o | or | o | or |
| C08_R | c | nc | f | f | r | r | or | r | or | r | or | r | or | r | or |
| C09_L | c | nc | f | f | r | r | or | r | or | r | or | r | or | r | or |
| C10_R | c | me | nc | f | o | o | or | o | or | o | or | o | or | o | or |
| C11_L | c | me | nc | f | o | o | or | o | or | o | or | o | or | o | or |
| C12_R | c | nc | t | t | o | o | or | o | or | o | or | o | or | o | or |
| C13_L | c | nc | t | t | o | o | or | o | or | o | or | o | or | o | or |
| C14_C | c | me | v | v | o | o | or | o | or | o | or | o | or | o | or |
| C15_R | c | me | v | v | v | v | v | v | v | vba | vba | v | v | vba | vba |
| C16_L | c | me | v | v | v | v | v | v | v | vba | vba | v | v | vba | vba |
| C17_C | c | me | v | v | v | v | v | v | v | vba | vba | v | v | vba | vba |
| C18_R | c | me | v | v | v | v | v | v | v | vba | vba | v | v | vba | vba |

| | | | | | | | | | | | | | | | |
|---|---|---|---|---|---|---|---|---|---|---|---|---|---|---|---|
| C19_L | c | me | v | v | v | v | v | v | v | vba | vba | v | v | vba | vba |
| C20_R | c | me | v | v | v | v | v | v | v | vba | vba | v | v | vba | vba |
| C21_L | c | me | v | v | v | v | v | v | v | vba | vba | v | v | vba | vba |
| C22_R | c | me | b | ba | b | b | b | ba | ba | vba | vba | ba | ba | vba | vba |
| C23_L | c | me | b | ba | b | b | b | ba | ba | vba | vba | ba | ba | vba | vba |
| C24_R | c | me | b | ba | b | b | b | ba | ba | vba | vba | ba | ba | vba | vba |
| C25_L | c | me | b | ba | b | b | b | ba | ba | vba | vba | ba | ba | vba | vba |
| C26_R | c | me | b | ba | b | b | b | ba | ba | vba | vba | ba | ba | vba | vba |
| C27_L | c | me | b | ba | b | b | b | ba | ba | vba | vba | ba | ba | vba | vba |
| C28_R | c | me | b | bu | b | b | b | bu | bu | bu | bu | bu | bu | bu | bu |
| C29_L | c | me | b | bu | b | b | b | bu | bu | bu | bu | bu | bu | bu | bu |
| C30_R | c | me | b | bu | b | b | b | bu | bu | bu | bu | bu | bu | bu | bu |
| C31_L | c | me | b | bu | b | b | b | bu | bu | bu | bu | bu | bu | bu | bu |
| C32_R | c | nc | f | f | r | r | or | r | or | r | or | r | or | r | or |
| C33_L | c | nc | f | f | r | r | or | r | or | r | or | r | or | r | or |
| C34_R | c | nc | f | f | r | m | m | r | or | r | or | m | m | m | m |
| C35_L | c | nc | f | f | r | m | m | r | or | r | or | m | m | m | m |
| C36_R | c | nc | f | f | m | m | m | m | m | m | m | m | m | m | m |
| C37_L | c | nc | f | f | m | m | m | m | m | m | m | m | m | m | m |

Supplementary S4. Top hypotheses of modularity across the Sminthopsin cranium determined several data filtering pipelines (row-column combinations). All model selections were performed using subsampleEMMLi() in EMMLi2 (Goswami et al., 2019; Goswami & Finarelli, 2016). Datasets were subsampled 100 times to 80% of the total landmark set (274 for all, 95 for only point landmarks) to ensure robust model selection. Only four models were identified as the best fit in any of the analyses: Goswami (bullae as separete, premolar as molar, sep between) (G_BuS_PmM_Sep), Goswami (bullae as basicranium, premolar as molar, sep between) (G_BuB_PmM_Same), Goswami (bullae as separate, premolar as molar, sep between) (G_BuS_PmM_Sep), Goswami (bullae as separate, premolar as rostrum, sep between) (G_BuS_PmR_Sep). Number in brackets denotes the number of subsamples out of 100 which supported this model, those above 95 are shown in bold.

| Genera | GPA | Landmarks | No correction | Species variation removed | Allometry corrected (shared slope) |
|---|---|---|---|---|---|
| All | Single | All | G_BuS_PmM_Sep (74) | G_BuS_PmM_Sep (75) | G_BuS_PmM_Sep (68) |
| All | Single | Points only | **G_BuS_PmM_Sep (97)** | **G_BuS_PmM_Sep (100)** | G_BuS_PmM_Sep (93) |
| All | Multi. | All | G_BuS_PmM_Sep (86) | **G_BuS_PmM_Sep (100)** | NA |
| All | Multi. | Points only | G_BuS_PmM_Sep (70) | G_BuS_PmM_Sep (95) | NA |
| *Sminthopsis macroura clade* | Single | All | **G_BuS_PmR_Sep (98)** | G_BuS_PmR_Sep (90) | G_BuS_PmR_Sep (84) |
| *Sminthopsis macroura clade* | Single | Points only | G_BuS_PmM_Sep (65) | **G_BuS_PmM_Sep (100)** | G_BuS_PmM_Sep (72) |
| *Sminthopsis macroura clade* | Multi. | All | **G_BuS_PmM_Sep (100)** | **G_BuS_PmM_Sep (100)** | NA |
| *Sminthopsis macroura clade* | Multi. | Points only | G_BuS_PmM_Sep (73) | **G_BuS_PmM_Sep (97)** | NA |
| *Ningaui* | Single | All | **G_BuS_PmM_Sep (100)** | **G_BuS_PmM_Sep (100)** | **G_BuS_PmM_Sep (100)** |
| *Ningaui* | Single | Points only | G_BuS_PmM_Sep (26) | G_BuB_PmM_Sep (80) | G_BuS_PmM_Sep (51) |
| *Ningaui* | Multi. | All | **G_BuS_PmM_Sep (100)** | G_BuS_PmM_Sep (91) | NA |
| *Ningaui* | Multi. | Points only | G_BuS_PmM_Sep (54) | G_BuS_PmM_Sep (80) | NA |
| *Sminthopsis murina clade* | Single | All | **G_BuS_PmM_Sep (99)** | **G_BuS_PmM_Sep (99)** | **G_BuS_PmM_Sep (96)** |
| *Sminthopsis murina clade* | Single | Points only | **G_BuS_PmM_Sep (100)** | G_**BuS_PmM_Sep (97)** | **G_BuS_PmM_Sep (100)** |
| *Sminthopsis murina clade* | Multi. | All | **G_BuS_PmM_Sep (100)** | G_BuS_PmM_Sep (92) | NA |
| *Sminthopsis murina clade* | Multi. | Points only | G_BuS_PmM_Sep (54) | G_BuS_PmM_Sep (64) | NA |
| *Antechinomys* | Single | All | G_BuS_PmM_Sep (91) | G_BuS_PmM_Sep (91) | G_BuS_PmM_Sep (93) |
| *Antechinomys* | Single | Points only | G_BuS_PmM_Sep (93) | **G_BuS_PmM_Sep (97)** | **G_BuS_PmM_Sep (99)** |
| *Antechinomys* | Multi. | All | **G_BuS_PmM_Sep (100)** | **G_BuS_PmM_Sep (96)** | NA |
| *Antechinomys* | Multi. | Points only | G_BuB_PmM_Sep (51) | G_BuS_PmM_Sep (88) | NA |

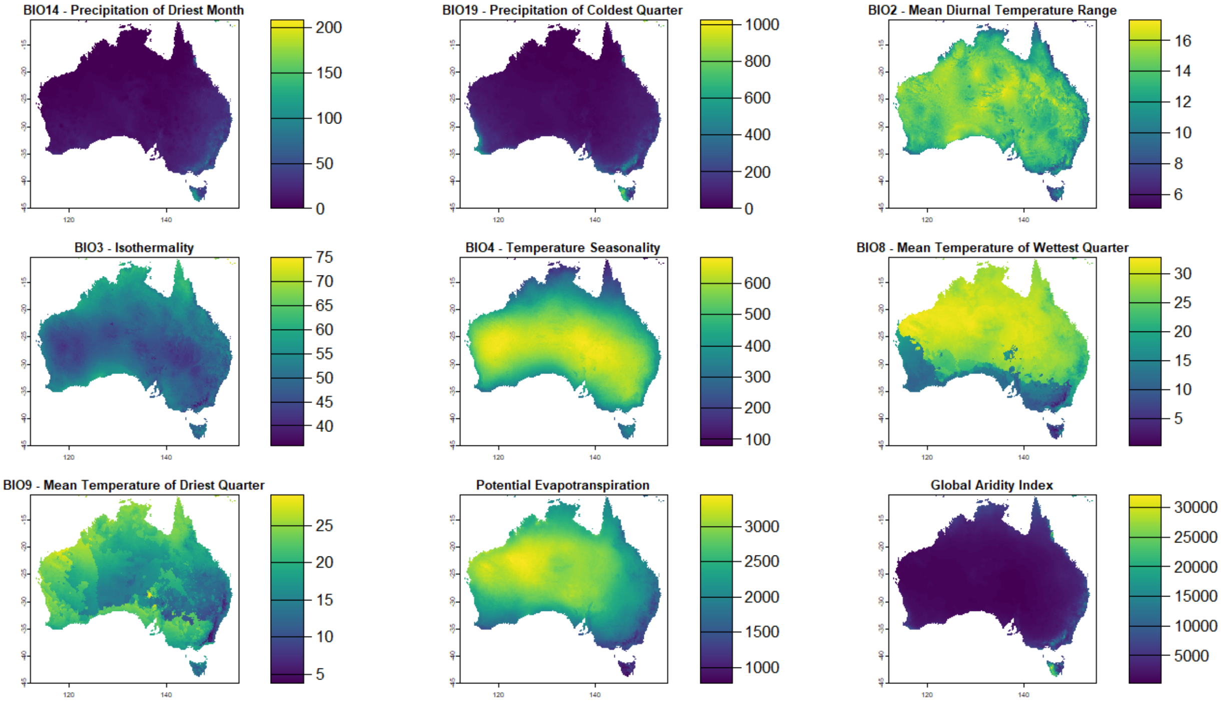

Supplementary S5. Maps of environmental variables used in generalized dissimilarity modelling of cranial module dissimilarity against environmental dissimilarity. A-G) WorldClim variables downloaded from https://www.worldclim.org/data/worldclim21.html (Fick & Hijmans, 2017). H-I) Potential evapotranspiration and global aridity index downloaded from https://doi.org/10.6084/m9.figshare.7504448.v5 (Zomer et al., 2022).

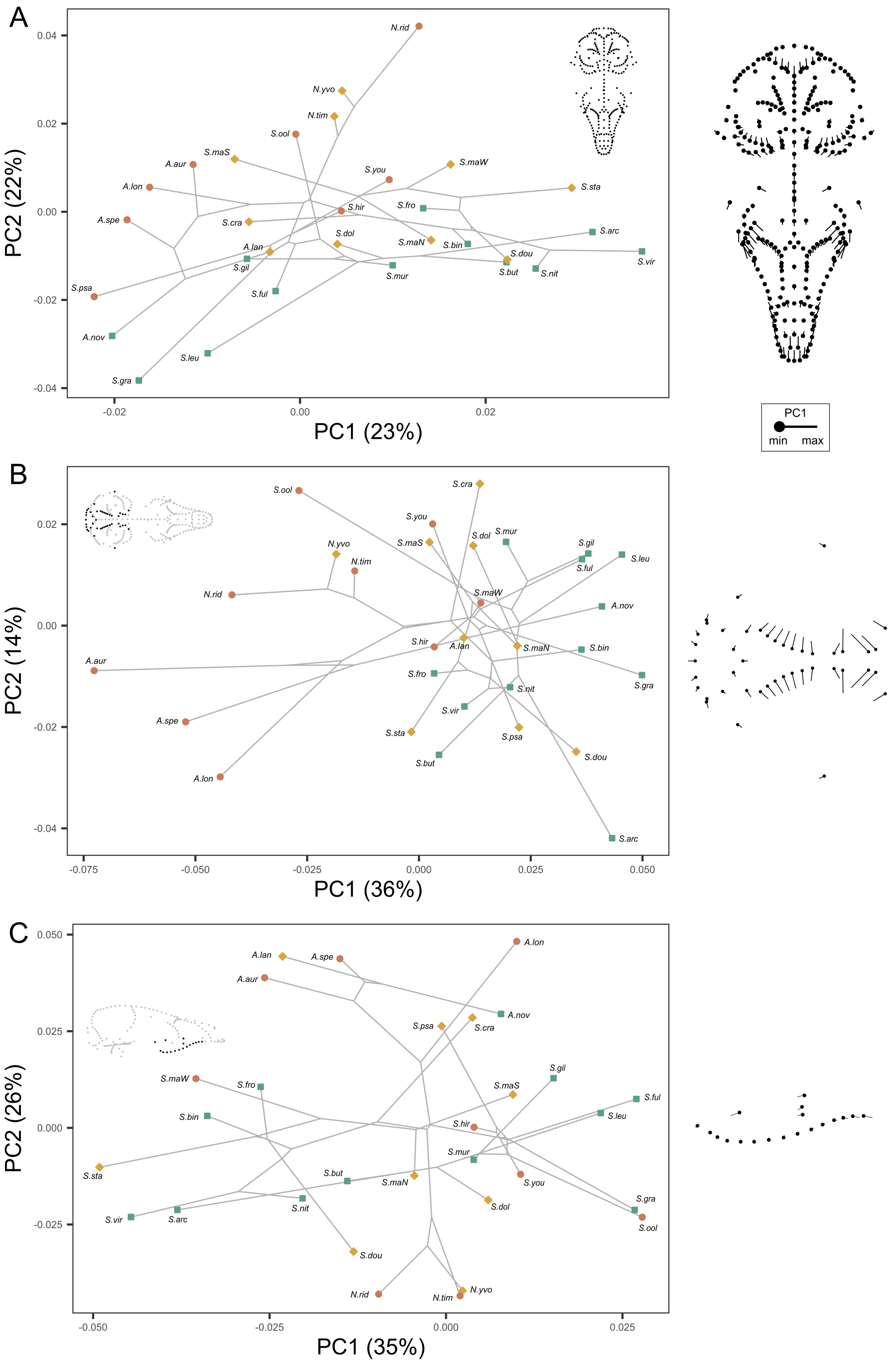

A
PC2 (22%)
PC1 (23%)
0.04
0.02
0.00
-0.02
-0.04
-0.02
0.00
0.02
N.rid
N.yvo
N.tim
S.ool
S.maS
A.aur
S.maW
S.you
A.lon
S.sta
S.fro
S.hir
A.spe
S.cra
S.arc
S.dol
S.maN
S.bin
A.lan
S.vir
S.dou
S.gil
S.mur
S.but
S.nit
S.ful
S.psa
A.nov
S.leu
S.gra
PC1
min
max
B
PC2 (14%)
PC1 (36%)
0.02
0.00
-0.02
-0.04
-0.075
-0.050
-0.025
0.000
0.025
0.050
S.ool
S.cra
S.you
S.maS
S.dol
S.mur
N.yvo
S.gil
N.tim
S.leu
S.ful
N.rid
S.maW
A.nov
A.aur
S.hir
A.lan
S.maN
S.bin
S.fro
S.gra
S.nit
S.vir
A.spe
S.sta
S.psa
S.dou
S.but
A.lon
S.arc
C
PC2 (26%)
PC1 (35%)
0.050
0.025
0.000
-0.025
-0.050
-0.025
0.000
0.025
A.lan
A.spe
A.lon
A.aur
A.nov
S.psa
S.cra
S.maW
S.fro
S.gil
S.maS
S.bin
S.ful
S.leu
S.hir
S.mur
S.sta
S.but
S.maN
S.you
S.dol
S.gra
S.vir
S.arc
S.nit
S.ool
S.dou
N.rid
N.tim
N.yvo

Supplementary S6. Phylogenetic PCA (Revell, 2009) of skull shape within the Sminthopsinae by module: A) whole skull, B), basicranium, C) molar. Species are colored by environment. Analyses were carried out using the mean shape for each species following separate module-specific GPAs. Displacement vector plots show the changes in shape from the minimum observed PC1 value (solid circle) to the maximum observed PC1 value (end of vector) for each module. Plots of the other modules can be found in Figure 2. Similar plots for all modules generated using a single whole-skull GPA and allometry corrected data can be found in Supplementary S7-8 respectively.

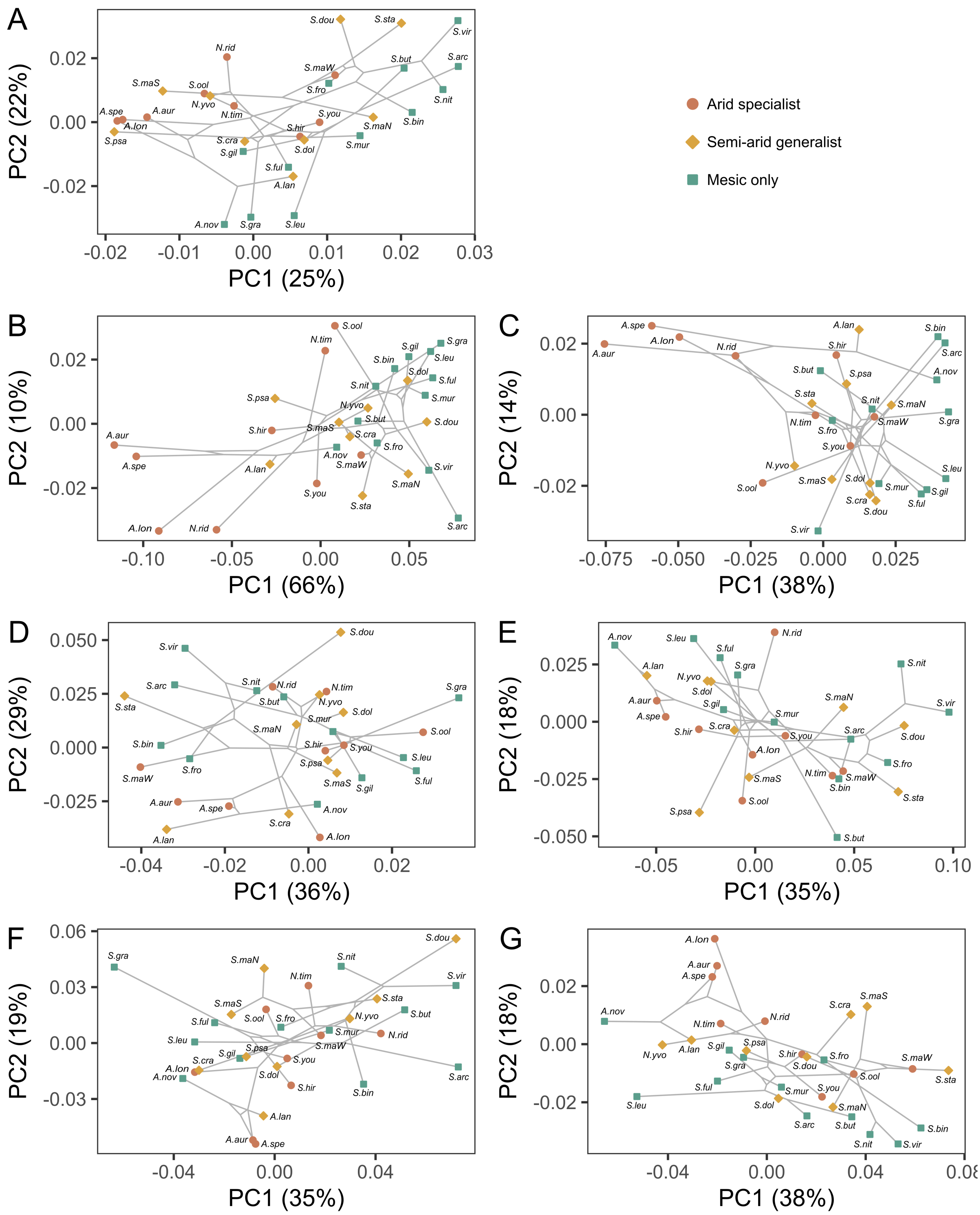

Supplementary S7. PhyloPCA (Revell, 2009) across the Sminthopsinae cranium by module with allometric shape removed: A) whole cranium, B) bullae, C) basicranium, D) molar, E) orbital, F) rostrum, G) vault. PhyloPCA components were generated using the mean shape for each species following separate module-specific GPAs. Allometric shape was extracted separately for each module using whole-skull centroid size.

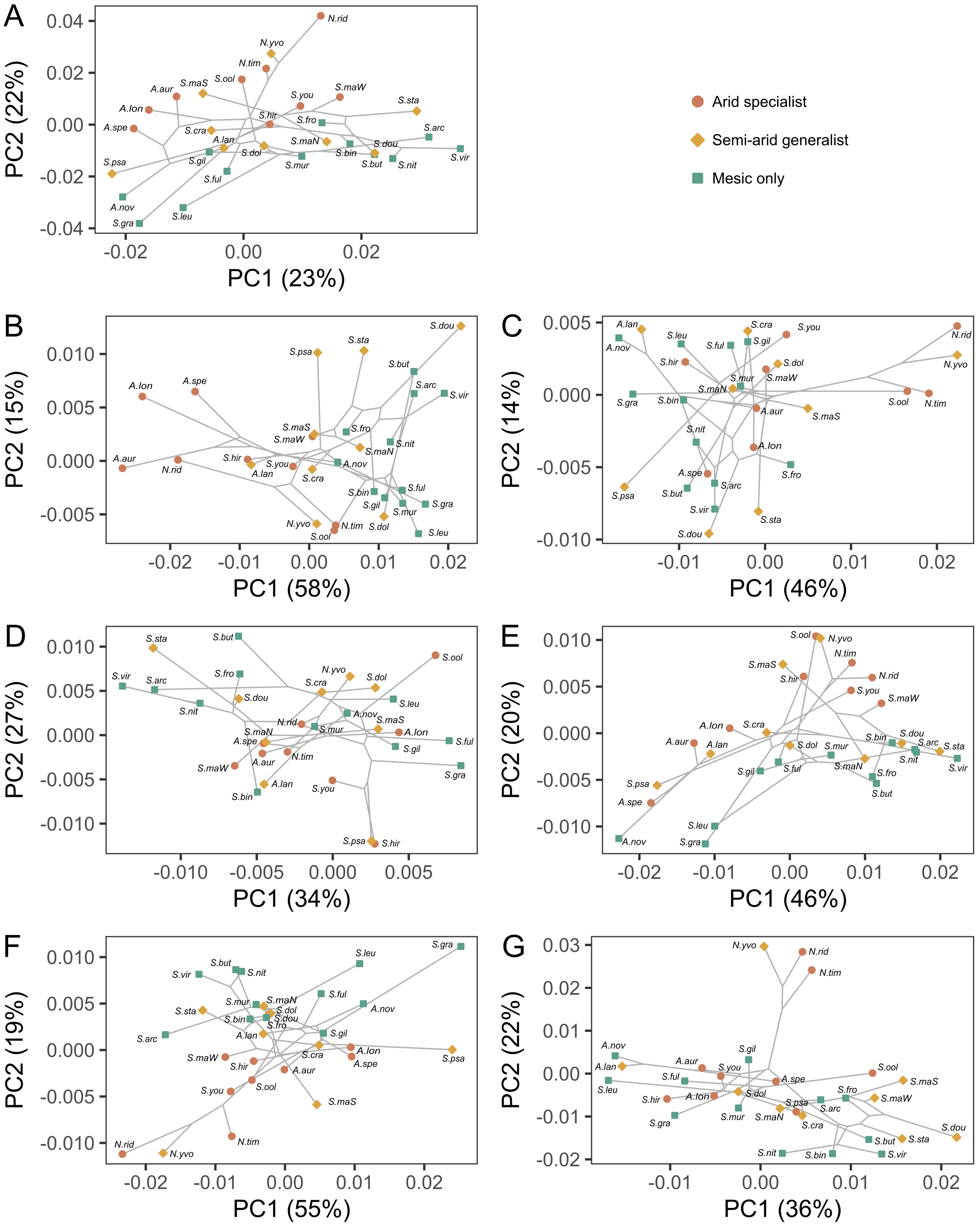

Supplementary S8. PhyloPCA (Revell, 2008) within the Sminthopsinae cranium by module using a single Procrustes superimposition for the entire cranium: A) whole cranium, B) bullae, C) basicranium, D) molar, E) orbital, F) rostrum, G) vault. PhyloPCA components were generated using the mean shape for each species.

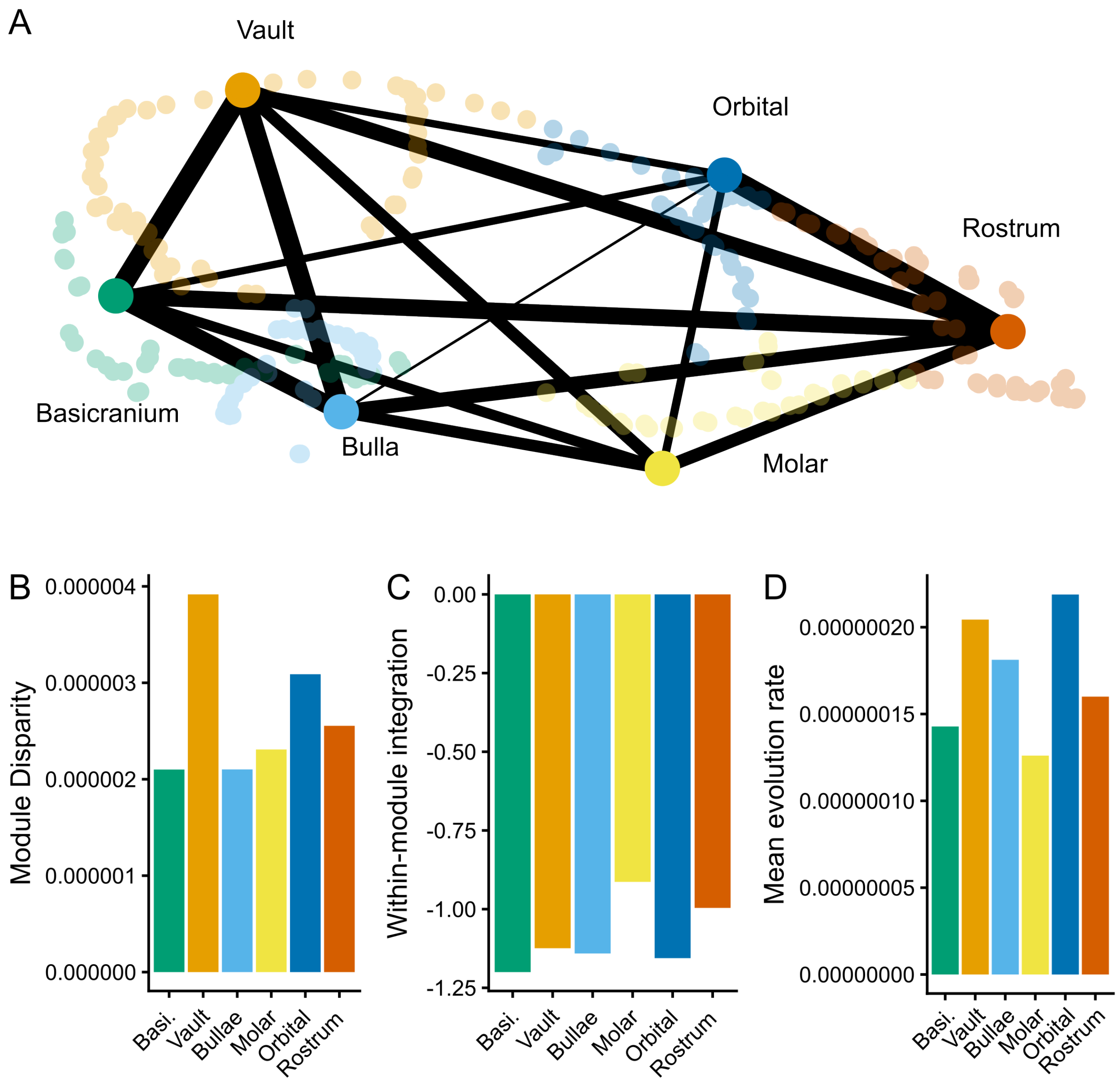

**Supplementary S9.** Patterns of disparity and integration between modules across the Sminthopsin cranium using a single whole-skull GPA. (A) Relative between-module integration, width of edges represents relative covariance between landmarks of each module pair, r values range from 0.408 (bullae:vault) to 0.708 (bullae: basicranium). (B) Total disparity (Procrustes variance) of each module. (C) Within-module integration, lower (more negative) values indicate weaker integration. Lines joined to bars indicate significant differences between integration values (D) Evolution rate of each module under a Brownian Motion model of evolution. Pairwise significant differences between measures are shown as ticks on the line joined to the focal module.

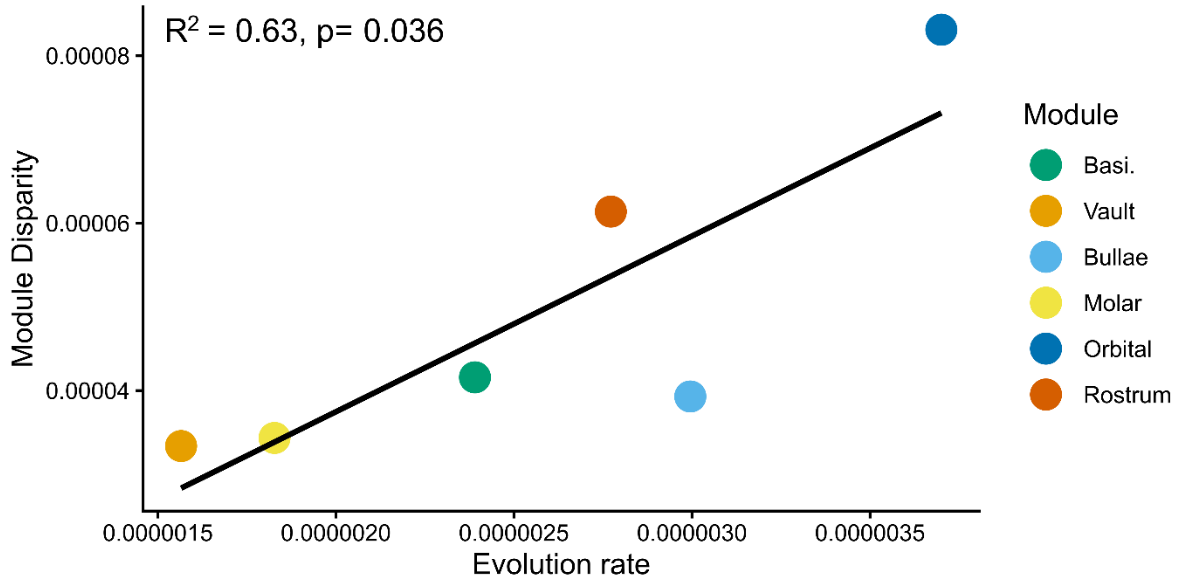

**Supplementary S10.** Relationship between module disparity and mean evolution rate calculated assuming Brownian Motion.

Supplementary S11. Sum of i-spline coefficients for each significant predictor of morphological dissimilarity for the entire cranium and each module separately following generalised dissimilarity modelling (GDM).

| dataset | % variance explained | Genetic distance | Skull size | Latitude | Isothermality | Temperature Seasonality | Aridity Index | Mean diurnal temperature range | Longitude | Precipitation of driest month | Mean temperature of wettest quarter | Precipitation of coldest quarter | Potential evapotranspiration | Geographic distance | Mean temperature of driest quarter |
|---|---|---|---|---|---|---|---|---|---|---|---|---|---|---|---|
| whole | 33.966 | 0.026 | 0.013 | 0.006 | 0.01 | 0.006 | | | 0.001 | | 0.002 | 0.002 | | | |
| bulla | 32.053 | 0.035 | 0.021 | | | 0.06 | 0.015 | 0.012 | 0.007 | | 0.007 | | 0.009 | | |
| basicranium | 25.731 | 0.028 | 0.014 | | | 0.03 | 0.008 | 0.002 | 0.001 | | 0.004 | 0.006 | 0.004 | | |
| molar | 16.298 | 0.025 | 0.007 | 0.013 | 0.012 | 0.003 | | 0.002 | 0.001 | 0.008 | 0.001 | 0.001 | | | |
| orbital | 18.258 | 0.033 | 0.014 | 0.029 | 0.016 | | | 0.005 | 0.001 | 0.015 | | | | 0.005 | |
| rostrum | 14.643 | 0.035 | 0.018 | 0.012 | 0.01 | 0.014 | 0.006 | | | | 0.002 | | | 0.002 | |
| vault | 19.43 | 0.035 | 0.014 | 0.017 | 0.009 | 0.003 | 0.001 | 0.002 | 0.003 | | 0.005 | 0.005 | | | 0.009 |

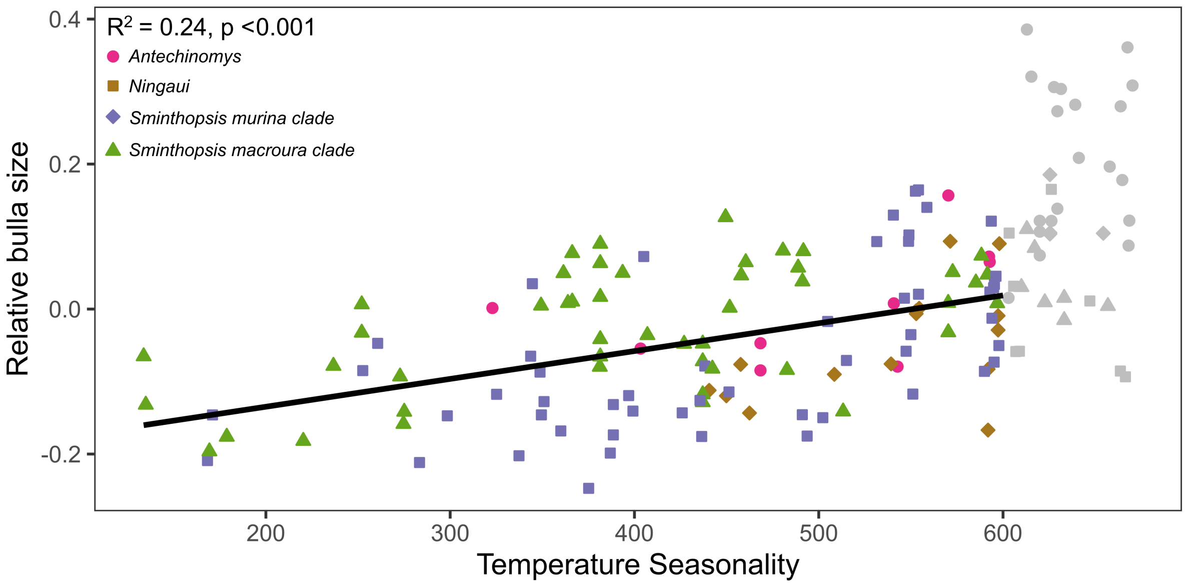

Supplementary S12. Post-hoc comparison of the effect of Temperature Seasonality (BIO4) (Fick & Hijmans, 2017) on relative bullae size following a Generalized Dissimilarity Model (GDM) identifying Temperature Seasonality as a strong predictor of bullae shape (see Figure 4). Relative bulla size was calculated as the residuals of a regression between bulla centroid size and 'rest of the skull' centroid size. This plot shows the overall pattern following the removal of samples from areas of extreme aridity (Temperature seasonality >600), excluded points are shown in grey.

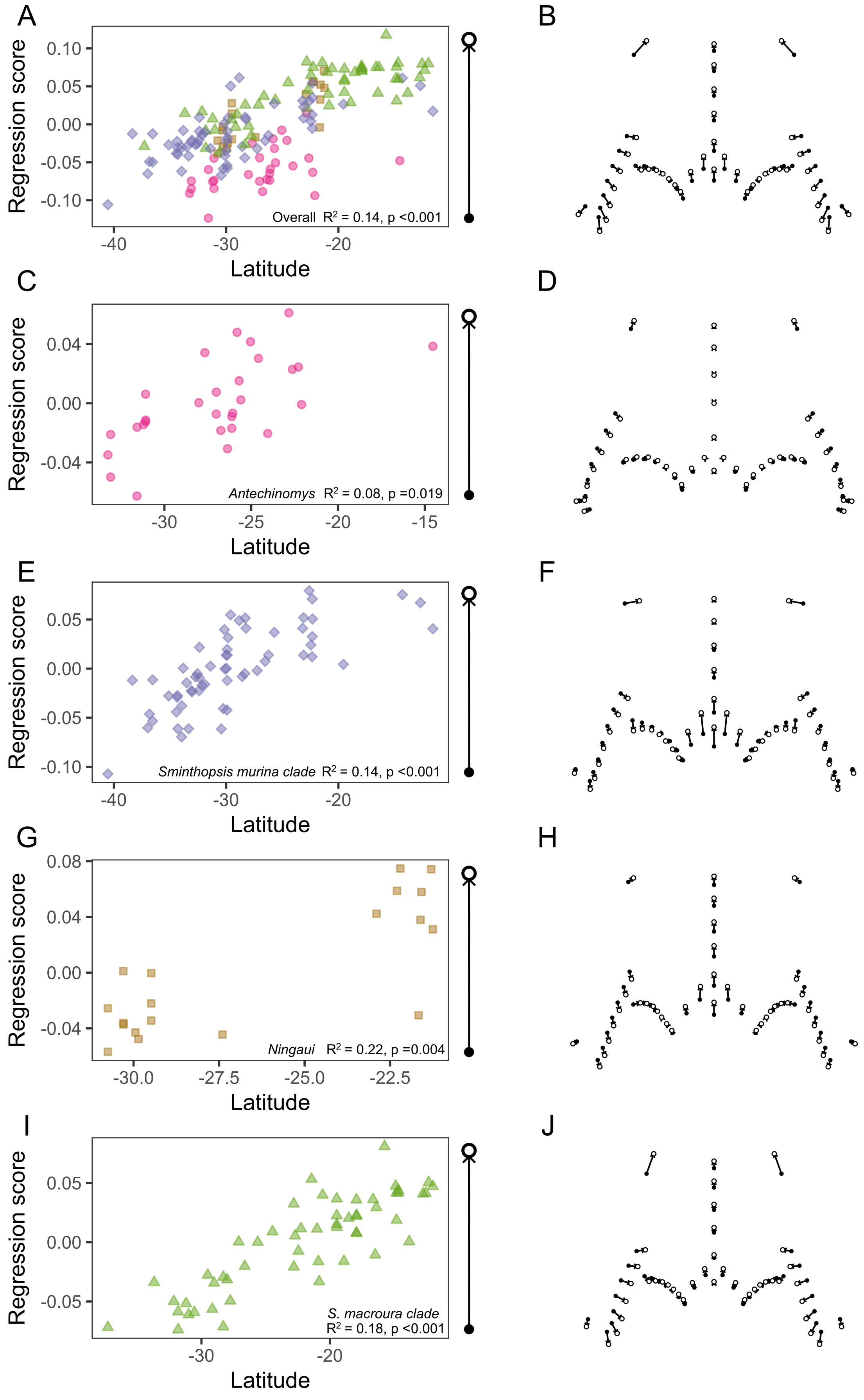

Supplementary S13. Orbital shape associated with changes in Latitude following a Procrustes

ANOVA and linear regression (procD.lm). Performed for all data and for each genus individually. A, C, E, G, and I (right) show regression score ~ latitude for each analysis. B, D, F, H, and J show displacement vectors (arrows) of orbital shape from the lowest regression score (filled circle) to the highest regression score (filled circle) for each model. Note that all analyses have displacement vectors which show a narrowing of the inter-orbital region as latitude increases.
